# Scaffold-Lab: Critical Evaluation and Ranking of Protein Backbone Generation Methods in A Unified Framework

**DOI:** 10.1101/2024.02.10.579743

**Authors:** Zhuoqi Zheng, Bo Zhang, Bozitao Zhong, Kexin Liu, Zhengxin Li, Junjie Zhu, Jinyu Yu, Ting Wei, Hai-Feng Chen

## Abstract

*De novo* protein design has undergone a rapid development in recent years, especially for backbone generation, which stands out as more challenging yet valuable, offering the ability to design novel protein folds with fewer constraints. However, a comprehensive delineation of its potential for practical application in protein engineering remains lacking, as does a standardized evaluation framework to accurately assess the diverse methodologies within this field. Here, we proposed Scaffold-Lab benchmark focusing on evaluating unconditional generation across metrics like designability, novelty, diversity, efficiency and structural properties. We also extrapolated our benchmark to include the motif-scaffolding problem, demonstrating the utility of these conditional generation models. Our findings reveal that *FrameFlow* and *RFdiffusion* in unconditional generation along with *Rfdiffusion* and GPDL in conditional generation showcased the most outstanding performances. Furthermore, we described a systematic study to investigate conditional generation and applied it to the motif-scaffolding task, offering a novel perspective for the analysis and development of conditional protein design methods. All data and scripts will be available at https://github.com/Immortals-33/Scaffold-Lab.

## Introduction

Proteins are fundamental to biological systems due to their diverse functions, largely attributed to the hierarchical organization that progresses from amino acid sequences to three-dimensional (3D) structures^1,2^. Despite substantial kinds of proteins already exist in nature, they have just covered a small fraction of the whole potential protein sequence space^3^. To this extent, *computational protein design* has been developed to extend the functionality of proteins. The magnificent breakthroughs in protein design along with protein structure prediction^4^ have allowed biologists to dive deeper into the mystery of protein structures, breathing a new life into the field of protein science and protein engineering^5,6^.

Protein design has undergone a paradigm shift over the past few decades. In the early stages of *de novo* protein design, researchers used physical-based methods to search through rotamer libraries in order to design new protein sequences based on a pre-defined structure scaffold^7^. In the past couple of years, the rapid development of deep learning has greatly invigorated the field of computational structural biology, both in protein sequence design^8–15^ and protein structure prediction^4,16–21^. Although given the vast sequence space to be explored^3^, it has been long acknowledged that proteins realize their biological functions directly by 3D structures^1,22^. Protein backbone generation was carried out to directly manipulate the topologies of proteins^23,24^, aiming to treat the 3D structure of protein as the design objective. While accurate modeling the structural constraints in 3D cartesian space is challenging for physical-based methods, the rapid advancements in generative models over the recent years, for example, denoising diffusion models^25,26^ have been a game changer with their exceptional performance in modeling high-dimensional data distribution^27^, which has already achieved incremental progress in several works^28–34^.

Although protein backbone generation has seen a surge of interest in protein design, several limitations have yet to be solved. First of all, most of those methods use a relatively “coarse-grained” way to either treat the all-atom protein structure as a backbone-only or *C*_*⍺*_-only protein. This compression leads to the result that only the information of protein backbone, or namely a “scaffold” can be generated through these methods, where a sequence design method is needed later to find the optimal sequences that folding into the designed backbones. Moreover, since protein backbone generation is targeted for novel structure generation and is an upstream step in protein engineering, there is still a lack of knowledge about how these methods can be applied to real-world applications. Furthermore, due to the varying metrics and length scopes employed by different methods, a set of benchmark for assessing different protein backbone generation methods is urgently needed, especially compared to the various established works in place for sequence design^35–38^, protein fitness prediction^39^ and molecular generation tasks^40^.

Here we proposed a unified benchmark, Scaffold-Lab, to evaluate current protein backbone generation methods extensively. We selected seven representative protein backbone generation methods based on their performance and availability, then carefully grouped them into different categories according to different lengths of proteins. We divided the benchmark sets into two different problems. The first one, namely *unconditional generation*, is the task that generates complete protein scaffolds under no restrictions, with user-specified protein length as the only pivotal input parameter. Unconditional generation has been a proof-of-concept exploration for almost all protein backbone generation methods as it implies the capability of how well those models learn the distribution of realistic protein structures, which is essential to assessing the methods’ performances. Opposite to the first one, *conditional generation* creates proteins under certain user-specified restrictions. Though it has not been explored to a vast extent, it is an important task since it is the destination to fully-programmed roadmap of protein design, aiming to develop *de novo* proteins with properties specified by scientists. We selected the most widely-used *motif-scaffolding* task as a representative under this category, whose goal is to generate novel scaffolds around one or multiple motifs extracted from native protein structures, which usually plays a key role in specific biological functions. This strategy is particularly useful in applications like designing novel enzymes^41^, binders^34,42,43^ or vaccines^44^ and has been widely explored and evaluated within the last few years^33,34,45–48^. Within this problem set, several computational metrics designed to evaluate these methods were recast into this evaluation framework followed by a ranking stage performed to assess the capability of different methods.

Beyond prevalent computational metrics, we also set up a personalized scheme to gain a deeper understanding of the progress and limitations of these methods. We first performed analysis on structural properties for unconditional generation, indicating that current methods exhibit a strong propensity for generating highly structured proteins. In the context of conditional generation, we transcended widely-used computational metrics and delved into the underlying reasons for different outcomes of designs. Furthermore, the capacity to create ideal protein structures is a prerequisite of successful designs rather than meeting certain conditional criteria. In summary, our research undertook a comprehensive assessment using a novel framework to elucidate the current state of progress and existing constraints within this domain, aiming to pave the way for its future development.

## Material and Methods

### Dataset

**Table 1** listed the methods and their corresponding tasks selected to be evaluated in Scaffold-Lab benchmark test. Both *Genie*^49^ and *FrameDiff*^50^ used the invariant point attention module brought by Jumper et al.^4^ as the key component of their architecture. *FrameDiff* utilized diffusion modeling in Riemannian manifolds to perform protein diffusion in a SE(3) space, while *RFdiffusion*^34^ integrated a diffusion models with a pretrained structure prediction model RoseTTAFold^19^ to greatly improve the performance. *Chroma* incorporated a diffusion model and a graph neural network (GNN) to generate protein structure and sequence simultaneously, with a scaling technique to improve the sampling speed to a large extent. Both *TDS*^46^ and *FrameFlow*^51^ was developed upon *FrameDiff*’s architecture, with the former used a particle filtering strategy to raise conditional sampling applied to motif-scaffolding task, and the latter replaced diffusion models with a flow-matching process to significantly elevate the efficiency and designability compared with previous works. Akin to the strategy in Wang et al.^45^, *GPDL*^48^ leveraged the single structure prediction method ESMFold^16^ combined with the constrained hallucination and inpainting strategies to generate highly designable and diverse scaffolds around given motifs.

**Table 1.**
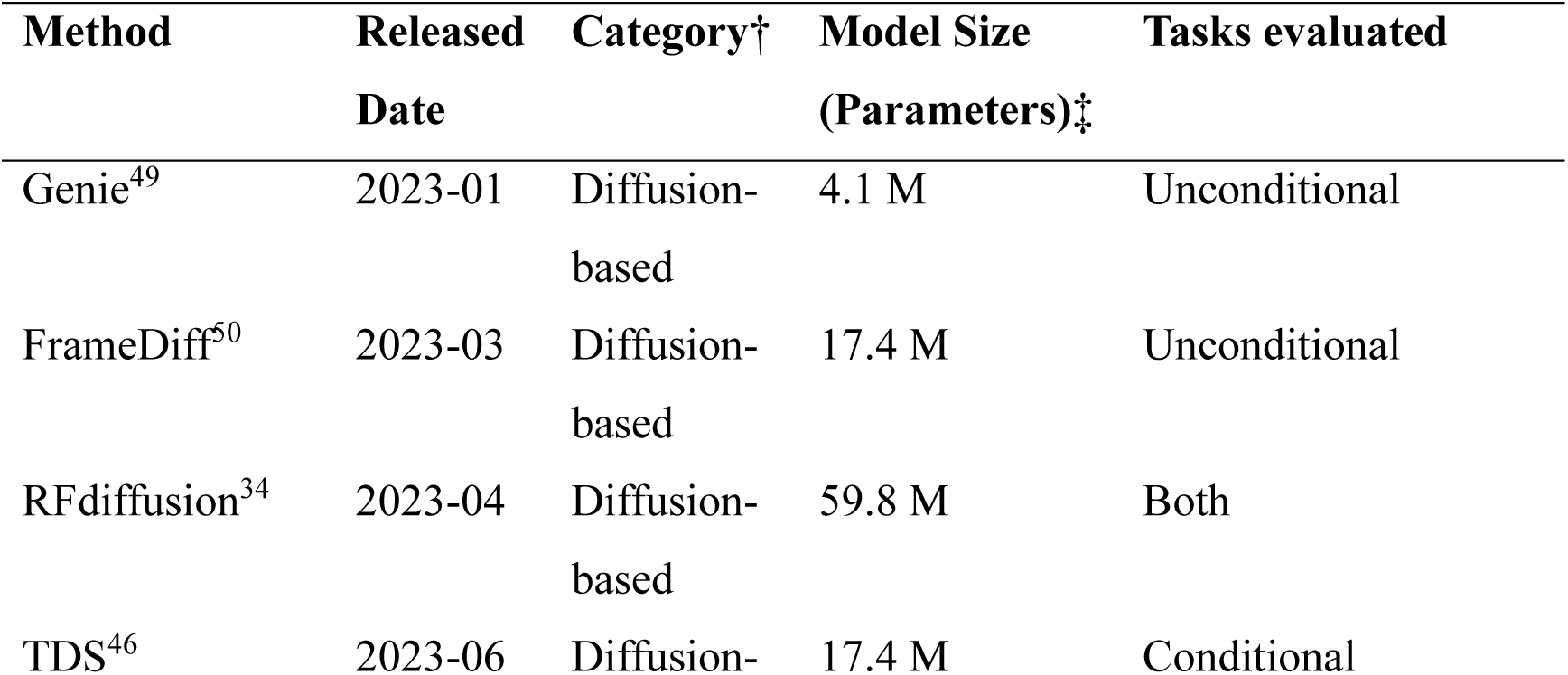
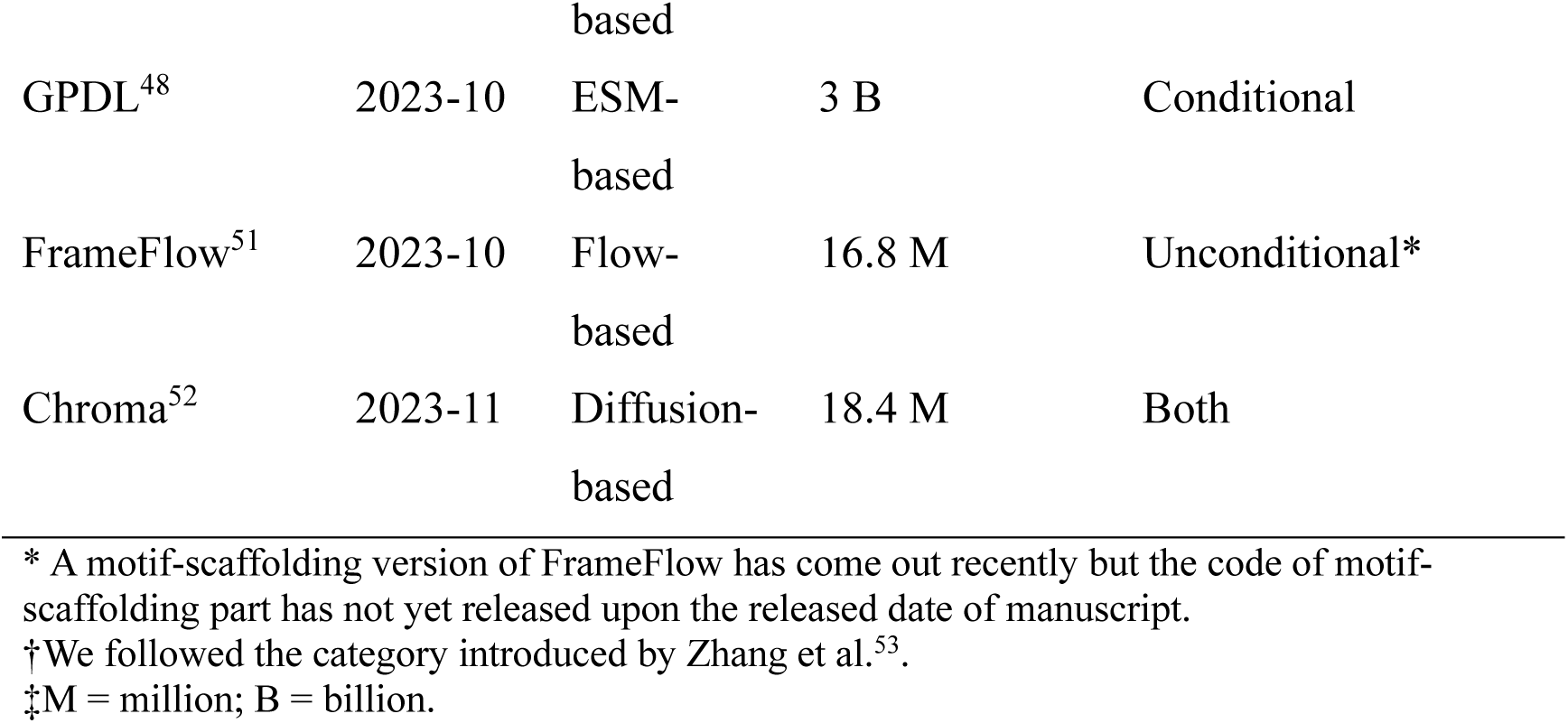
Methods selected to be tested in Scaffold-Lab.

Given the distinct application contexts and abilities to generate long sequences among various methods, it is crucial to categorize them appropriately prior to evaluation. Since the key parameter needed to be set by users in unconditional generation task is the desired length of generated proteins, we further grouped these methods into three sets (short, medium and long) based on the capable designed length scope of each method. After the grouping stage, some representative lengths were chosen within the scope, through which we seek an impartial and comparative evaluation for different methods within each set. **Table 2** lists the information after grouping by lengths. The analysis throughout this benchmark would be under this length-grouping framework.

**Table 2.** Length grouping of unconditional generation.

| Group | Length | Method |
| --- | --- | --- |
| Short | 50, 100 | Genie, FrameFlow, FrameDiff, Chroma, RFdiffusion |
| Medium | 200, 300, 400, 500 | FrameDiff, Chroma, RFdiffusion |
| Long | 600, 800, 1000 | Chroma, RFdiffusion |

For the motif-scaffolding task, we selected the dataset brought by *RFdiffusion*^34^. We excluded *6VW1*^54^ from our evaluation as it pertains to a multi-chain motif-scaffolding task which some of the selected methods were not equipped to design. Consequently, this resulted in a dataset comprising 24 cases. More detailed information is listed in **Supplementary Table 1**.

### Unconditional Generation

The overall workflow of Scaffold-Lab is shown in **Figure 1**. We performed unconditional protein backbone generation after grouping different methods by lengths. For each unique length within each method, we generated 100 backbones and fed them into the refolding pipeline. With 100 sequences designed and 10 sequences selected, we obtained a total number of 1000 refolded protein structures, which were mainly utilized to calculating designability. For the purpose of avoiding the sequence designer bias to the best degree, we only used refolded structures for the analysis on designability, while the novelty, diversity, efficiency and structural properties are calculated directly towards the protein backbones generated by tested methods.

**Figure 1.**
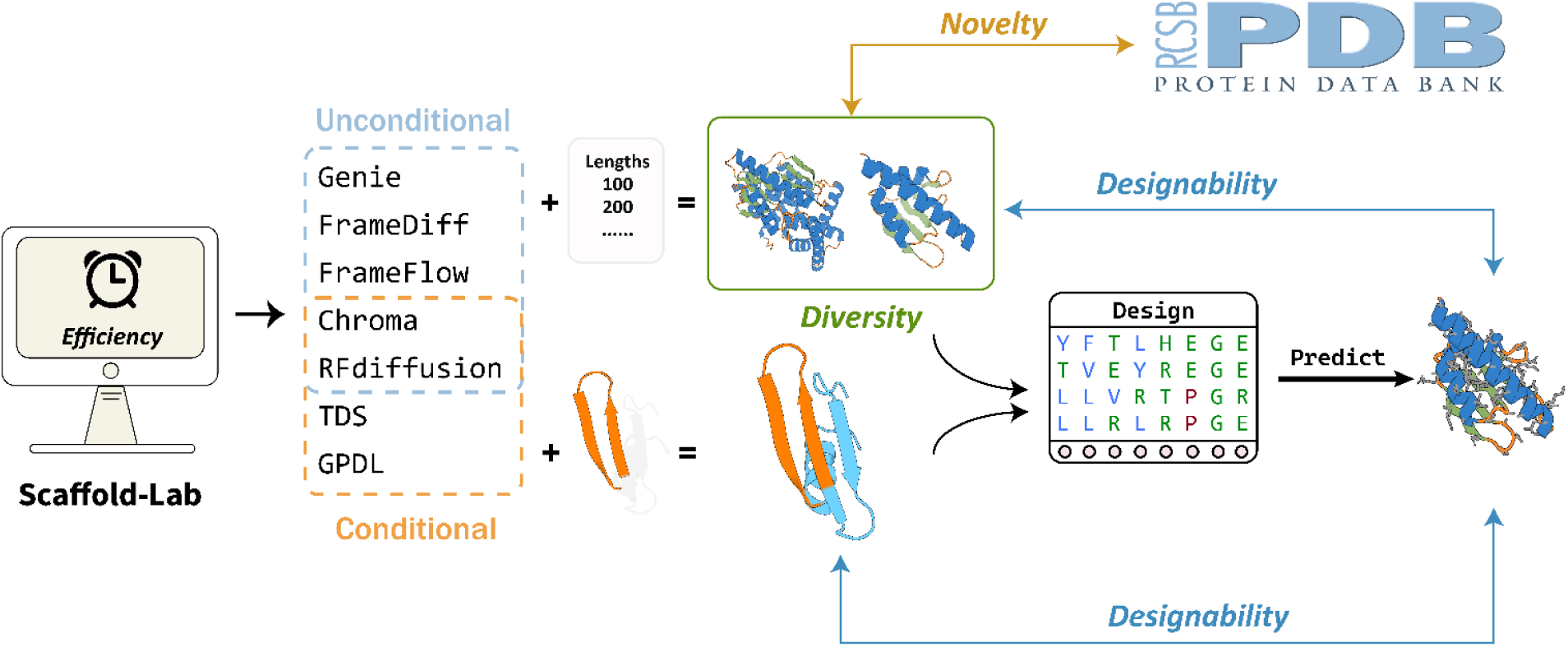
The overall workflow of Scaffold-Lab. Seven state-of-the-art protein backbone generation methods were evaluated by two types of tasks throughout a refolding pipeline. Four qualitative metrics were introduced to qualitatively assess the performances of methods, namely designability, novelty, diversity and efficiency.

We used ProteinMPNN^12^ as sequence designer throughout our pipeline which had been widely utilized in evaluation of protein backbone generation methods^34,49–51^ and validated by wet-lab experiments^34,45,55^. Out of consideration of the potential bias by different sequence design methods, we compared the performances ProteinMPNN to another state-of-the-art method, ESM-IF1^56^. The results in **Supplementary Figure 1** indicated that ProteinMPNN is able to find sequences with higher refolding ability than ESM-IF1, hence maximizing the potential of different protein backbone generation methods. For the structure predictor, we compared three prevalent highly accurate protein structure prediction methods, AlphaFold2^4^, ESMFold^16^ and OmegaFold^57^. The result showcased that three structure prediction methods exhibit a relatively strong correlation (shown in **Supplementary Figure 2)**. Thus, we used ESMFold out of its outstanding performance on *de novo* proteins and its high computational efficiency.

### Conditional Generation

For conditional generation, we chose *motif-scaffolding* as the essential problem to be solved in conditional generation, where the it is defined as below: Given a functional “motif” harbored in native proteins, these methods can generate “scaffold” around or between these motifs, where the lengths of scaffolds can sometimes be variable and randomly sampled. In our selected dataset, the motifs of native proteins, the length scope of scaffolds as well as scaffolding ways have been already defined, so we set up the evaluation experiments to use these methods to run through this dataset to generate plenty of protein backbones, which would enter the refolding pipeline to be examined the fidelities of different methods.

Since *TDS* and *GPDL* both need to specify the total length of protein, we designed backbones with mostly the same length within each case, where the length was chosen around the median value of length scope defined in Watson *et al*.^34^. For *Chroma* and *RFdiffusion* we performed length-variable design for each case. We randomly generated 10 sequences using ProteinMPNN and fixed the type of amino acids inside the motif regions during sequence design stage in order to stabilize the motif regions and improve the overall success rate.

### Evaluation Metrics

In this part, we introduced the metrics used in this evaluation, encompassing a range of both quantitative and qualitative aspects. We divided these metrics into five categories, namely *designability, novelty, diversity, efficiency* and *structural properties*, where the first four qualitative metrics would be used for ranking different methods. In the following sections, a brief introduction was done for each of these metrics.

### Designability

*Designability* measures the quality of generation, which is the most important metric for protein backbone generation tasks. Designability assesses the existence of any sequence capable to fold into the generated protein backbone. We used the self-consistency TM-score (*sc-TM*) metric raised by Trippe et al.^33^ where a protein backbone was defined as designable when the refolding structures and original backbones shares a TM-score^58^ > 0.5, which are considered to have the same fold pattern^59^. Compared to sc-RMSD, sc-TM could show better consistency through protein with different lengths (shown in **Supplementary Figures 3-4**). We also adopted a “Top *N* strategy”. While current works considered designing solely one sequence for the sc-TM criteria, the number of different sequences that fold into the generated structures could also reveal the robustness and fitness of backbones. Therefore, we reported the sc-TM as the mean value of *N* sequences with highest sc-TM among generated sequences, which defined as follows:

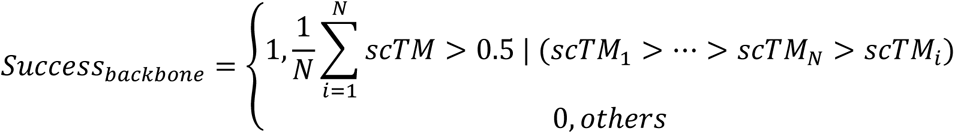

During our evaluation, we chose *N* = 3, which means we calculated the designability use the mean value of top 3 sequences with highest sc-TM values out of 10 selected sequences. We also reported the distribution of sc-TM and success rate by picking the best sequence as a fair comparison (shown in **Figure 2B** and **Supplementary Figure 5)**.

**Figure 2.**
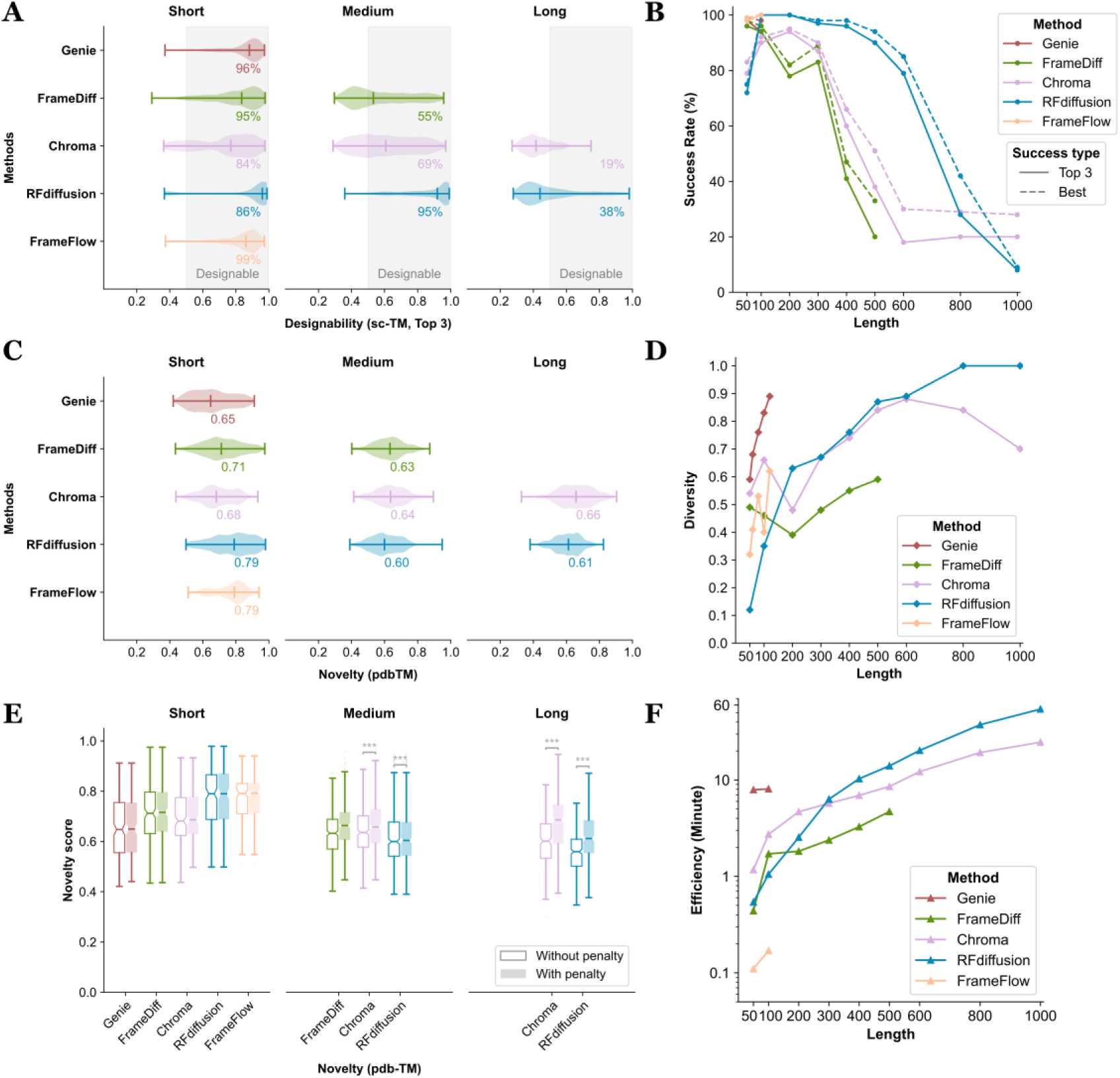
Evaluation results of unconditional generation. **(A)** Distribution of top 3 sc-TM. The number attached to each group denotes the proportion of backbones with top 3 sc-TM ≥ 0.5. **(B)** Success rates of tested methods. Success rate is defined by the number of backbones passing the sc-TM threshold divided by the total number of backbones within each group, with top 3 sc-TM by solid line and best sc-TM by dashed line. The text of numbers refers to the solid lines. **(C)** Distribution of pdb-TM values. The number attached to each group denotes the median value. **(D)** Diversity performances. Diversity is defined as the number of unique clusters divided by the total number of proteins within each group using Foldseek-Cluster. **(E)** Differences between novelty calculation by whether adding the penalty term of designability. Stars on each data group indicate the differences between two types of data within the group, calculated using the T-test. *, *p*-value < 0.05; **, *p*-value < 0.01; ***, *p*-value < 0.001. **(F)** Time usage for generating protein backbones among different methods. The efficiency is calculated as the average time usage per backbone for generating 100 backbones. All tests were conducted under identical hardware resources.

For unconditional generation, we defined a designable protein backbone as its top 3 sc-TM > 0.5. For conditional generation (motif-scaffolding), we separated the definition for backbone and motif: For backbones, we followed the criteria used in unconditional generation; for motifs, the refolded motif ought to have a motif-RMSD < 1.0 Å compared with the original input motif. The designability of a scaffolded backbone was defined as backbone plus motif designability.

### Novelty

*Novelty* measures the structural difference between the generated protein backbones to all existing proteins. We followed the novelty metric raised by Yim *et al*.^50^ by calculating the highest TM-score between generated protein backbones and all proteins in Protein Data Bank (PDB)^60,61^, which is referred as *pdb-TM*. If *pdb-TM* is low, it means that the generated backbones are novel compared to most of the existing experimentally solved native proteins, and vice versa. We used Foldseek^62^ to search generated backbones against the PDB database. to find their nearest protein structure. In order to make a fair comparison while maintaining computational efficiency, we applied the same parameter suite to all tested methods.

Apart from pdb-TM, we supposed that there exist certain generated backbones which are novel but undesignable. Thus, the *novelty score* was added a penalty to poorly designable backbones when calculating novelty, formulated as below:

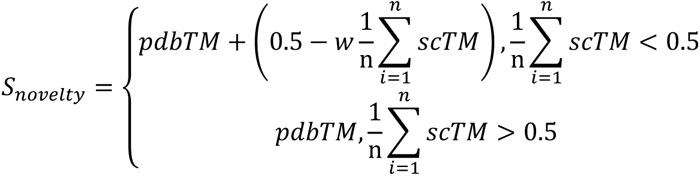

In the equation, *n* refers to the “Top N” strategy when picking sequences generated by ProteinMPNN, *w* means the degree of penalty, where higher *w* brings a more stringent penalty to the less-designable backbones. We set *n* = 3 and *w* = 1 throughout our experiments. Since we defined sc-TM > 0.5 as the threshold of designability, we left the backbones with sc-TM > 0.5 as original pdbTM value, otherwise a penalty is put on the shortfall part between sc-TM and the passing threshold, 0.5. After the reconsideration of novelty metric, the discrepancy between designable novel backbones and non-designable novel backbones would be exposed to a significant extent.

### Diversity

*Diversity* was designed to measure internal discrepancy of generated samples, reflecting the determinacy and the generalizability of models. Under the scenario of protein backbone generation, a method with greater diversity implies more diverse structural topologies can be selected as candidates for subsequent sequence design stage. This could benefit both the heterogeneity of potential application and the success rate for wet-lab experiments.

Since diversity is a referent metric of internal differences of generated examples, clustering the generated protein backbones is a natural way to assess the similarities between these protein backbones. We grouped the protein backbones by protein lengths and used Foldseek-Cluster^63^ to cluster each group of protein backbones. We used a clustering TM-Score threshold of 0.5 based on the aforementioned consensus^59^. For the definition of diversity, we followed Yim *et al*.^50^ as the number of unique clusters divided by the total number of protein backbones each group:

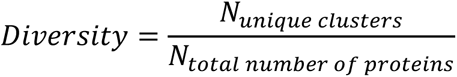

Generally, a higher count of clusters reflects more heterogeneous topologies appeared within a subset of sampled protein backbones, thus a higher diversity.

### Efficiency

Denoising diffusion models have ushered a new wave in the field of protein backbone generation as a representative of generative models. Despite their outstanding capabilities, many of them suffer from the iterative time-consuming denoising steps, which makes them suffer from the expensive consumption of both time and computational resources. Since efficiency is particularly crucial for the high-throughput scenario of protein engineering, we also compared the runtime of different protein backbone generation methods by generating 100 backbones per group and calculating the average time taken for each backbone. For the fairness of comparison, we maintained the hardware resources to be the same when performing experiments with different methods by conducting all experiments on a single CPU.

### Structural Properties

Proteins have various forms of internal structures organized in a hierarchical way. A protein with desired folded state usually has many unique covalent and non-covalent interactions which is a key theme in stabilizing the protein and leading it to an energy-favored state^2^. As protein backbone generation focus on the three-dimensional modeling of proteins, it is necessary to check the physical plausibility and quality from structural perspective. In this part, we analyzed the structural properties of protein from two different aspects, *secondary structure elements* and *radius of gyration*, respectively.

The secondary structure element (SSE) is a key feature of proteins since it directly represents the type of topologies and structural organization of a protein backbone. Since SSE is mainly determined by the backbone of protein and sequence design methods might bring internal bias for SSE preferences^37^, we directly computed SSE for the original generated protein backbones. We calculated SSE using DSSP software^64,65^ through Biotite^66^, count the eight types of SSEs assigned by DSSP and further group them into four common secondary structure types. We further analyzed the proportion of different SSEs in protein backbones generated by different groups to see if there’s any preference or potential rules produced by these methods.

Apart from SSEs, we also computed the radius of gyration (*Rg*) of each generated protein backbones. The radius of gyration of a protein can be measured in X-ray, neutron scattering and other experiments, which is one of the key properties of biological polymer molecule as it measures the size or compactness of proteins. *Rg* can be a key distinction reflect the flexibility difference between structured proteins and intrinsically disordered proteins (IDPs), which is of great significance to the function of proteins^67^. We computed *Rg* directly on the protein backbones generated by tested methods as what we applied in SSE computation, and compared to an experienced *Rg*-protein length diagram. The analysis on structural properties might provide a deeper insight into the protein backbones generated by current methods.

### Ranking

For the ranking of different methods, we used the widely utilized Technique for Order Preference by Similarity to Ideal Solution (TOPSIS). TOPSIS^68^ is a multi-criteria decision analysis method based on the concept that the shortest geometric distance from the positive ideal solution should be chosen as the best alternative, and vice versa. This method is usually used combined with comprehensive factor weighting methods to improve the confidence of evaluation, either subjective weighting methods like Analytic Hierarchy Process (AHP)^69^ or objective methods like CRiteria Importance Through Intercriteria Correlation (CRITIC)^70^. This ranking procedure, previously used in the evaluation of protein sequence design methods^37^, has been adapted for our work.

Although AHP performs well in many instances, it has been found the results based on AHP might significantly differ from the subjective weights^71^. Thus, we introduced CRITIC to objectively decide the importance of our metrics to minimize the impact of subjective factors on our ranking outcomes. After data preprocessing by setting values We used this data preprocessing procedure, and ranked different methods based on the length groups, where the four qualitative metrics are **designability**, **novelty**, **diversity** and **efficiency**. We suggested interested readers to Supplementary Text for a detailed version of our ranking procedure.

## Results

### Unconditional Generation

#### Designability

The overall results of unconditional generation task are shown in **Figure 2**. We first reported the top 3 sc-TM distribution of different methods along different groups and shown in **Figure 2A**. From the result of short-protein group, nearly all the evaluated methods performed well, where all the methods attained more than 85% designable backbones. Nevertheless, the performance of them dwindled as the lengths gradually increases, until neither the median sc-TM value of *RFdiffusion* nor *Chroma* reached the threshold of 0.5. We noted that *FrameFlow*, *Genie* and *FrameDiff* generated short proteins that almost all designable while *RFdiffusion* performed the most robust along all lengths.

To further explore the qualities of generated backbones, we calculated the success rates by both top 3 sc-TM and best sc-TM, whose results are shown in **Figure 2B**. From the trends displayed in the graph, we can observe that the success rate generally decreases as the length grows. We also noticed a watershed on the success rate: When the length is equal or less than 300, all of the tested methods show a strong performance with success rate around or more than 80%, which then rapidly dropped until less than 30% when it comes to proteins with not less than 800 residues. Additionally, the success rate between two distinct definitions indicates that the robustness of designed backbones also declines to a certain extent as the length increases.

Generally, for both the distribution of sc-TM and success rate, *RFdiffusion* performed the best among all tested methods, while other methods are also reliable when generating short proteins. When it comes to the situation of long proteins, it is essential to carefully check the plausibility of proteins before putting it into a downstream scene. The sc-RMSD values also supported the conclusion (**Supplementary Figure 4)**. As a reference, we also reported the best sc-TM distribution, which is shown in **Supplementary Figure 5**.

#### Novelty

We performed novelty analysis to different length groups, whose results are shown in **Figure 2C**. The pdb-TM values predominantly range between 0.6 and 0.7, with no significant differences in trends observed between various methods and length categories. Notably, *FrameFlow* and *RFdiffusion* exhibit slightly highest pdb-TM values in the group of short proteins, suggesting a tendency to generate protein backbones more akin to natural proteins. Given their relatively higher success rates in corresponding categories shown in Figure 2B, we hypothesized that this trend in novelty might also reflect the fitness of model to some extent, which implies a potential trade-off between novelty and designability.

To further validate our hypothesis, we used the novelty score with penalty put on protein backbones with low designability, in order to discover the correlation between these two methods. **Figure 2E** shows the differences between the distribution of pdb-TM and novelty score, where we conducted one-sided *Mann-Whitney U test*^72^ across different groups distinguished by with or without the penalty term. The results show that some of the groups have significant differences with *p-values* < 0.001, including *FrameDiff*, *Chroma* in medium-protein group (6.11 × 10^−8^ *and* 5.19 × 10^−4^, respectively) and *RFdiffusion, Chroma* in long-protein group ( 2.46 × 10^−1^^5^ *and* 3.33 × 10^−1^^6^ , respectively) . We also noticed that *RFdiffusion* and *FrameFlow* in the short protein group did not show significant differences in results with or without penalty, indicating that most proteins within those groups pass the designability threshold, which aligns with our previously proposed hypothesis. The differences between penalty groups grows along with proteins’ length, indicating the appearance of protein backbones with high novelty but low designability. Our analysis suggested that the designability should be partially taken into consideration when calculating novelty, whereas different methods did not show a strong variance in novelty overall. The complete results of significant test are listed in **Supplementary Table 2**.

#### Diversity

We conducted a diversity analysis upon different methods. We used the latest structure clustering algorithm Foldseek-Cluster to process the clustering step using a TM-score threshold of 0.5, where proteins with TM-score above this threshold would be clustered into the same group. The final diversity metric is defined as the number of unique clusters divided by the total number of proteins within each group, with a range from 0 to 1. The results are shown in **Figure 2D**.

*FrameFlow* had the lowest diversity compared with other methods, which probably owes to the diversity-designability trade-off mentioned in Yim *et al*.^50^. *RFdiffusion* and *Genie* showed a continuously increasing diversity by length within their length scope, where *RFdiffusion* started on a low point and reach almost a complete diverse performance when the length of protein is above 800. Interestingly, we also noticed that *FrameDiff* and *Chroma* shows a non-monotonic behavior, with *FrameDiff* declining at the beginning and climbing up later, while *Chroma* behaves in contrast. Generally, the diversity turns higher as the length of protein increases, where the redundancy of training set at different length levels can be a potential contributor.

#### Efficiency

The result displayed in **Figure 2F** demonstrate significant differences in efficiency among various methods. The tested methods showcase most variances when generating short proteins, as *Genie* took about 8 minutes whereas *FrameFlow* took just around 0.15 minutes to generate a backbone, where the latter exhibiting potential for high-throughput applications in conjunction with downstream uses. We also noticed that the growing trends of *Chroma* stabilized when generating long proteins which benefit from their scaling technique^52^, which is of great help when generating huge proteins. It is noteworthy that the speeds of current methods are still at the scale of minutes, remains a gap between advanced sequence design methods that generating sequences within a few seconds. Improving inference speed and modeling efficiency may represent an important direction for future development in this field.

### Analysis of Structural Properties

#### Secondary structure analysis

**Figure 3A** illustrates the SSEs composition from the perspectives of overall average. The average frequency of SSEs by different methods are all more inclined to generate structured proteins, with a *⍺*-helix percentage > 50% among most cases. We also noticed that this structured-preference did not change a lot even with the length increased up to 1000. The prefilter step during training where most loop-abundant structures were excluded might be key factor to this behavior.

**Figure 3.**
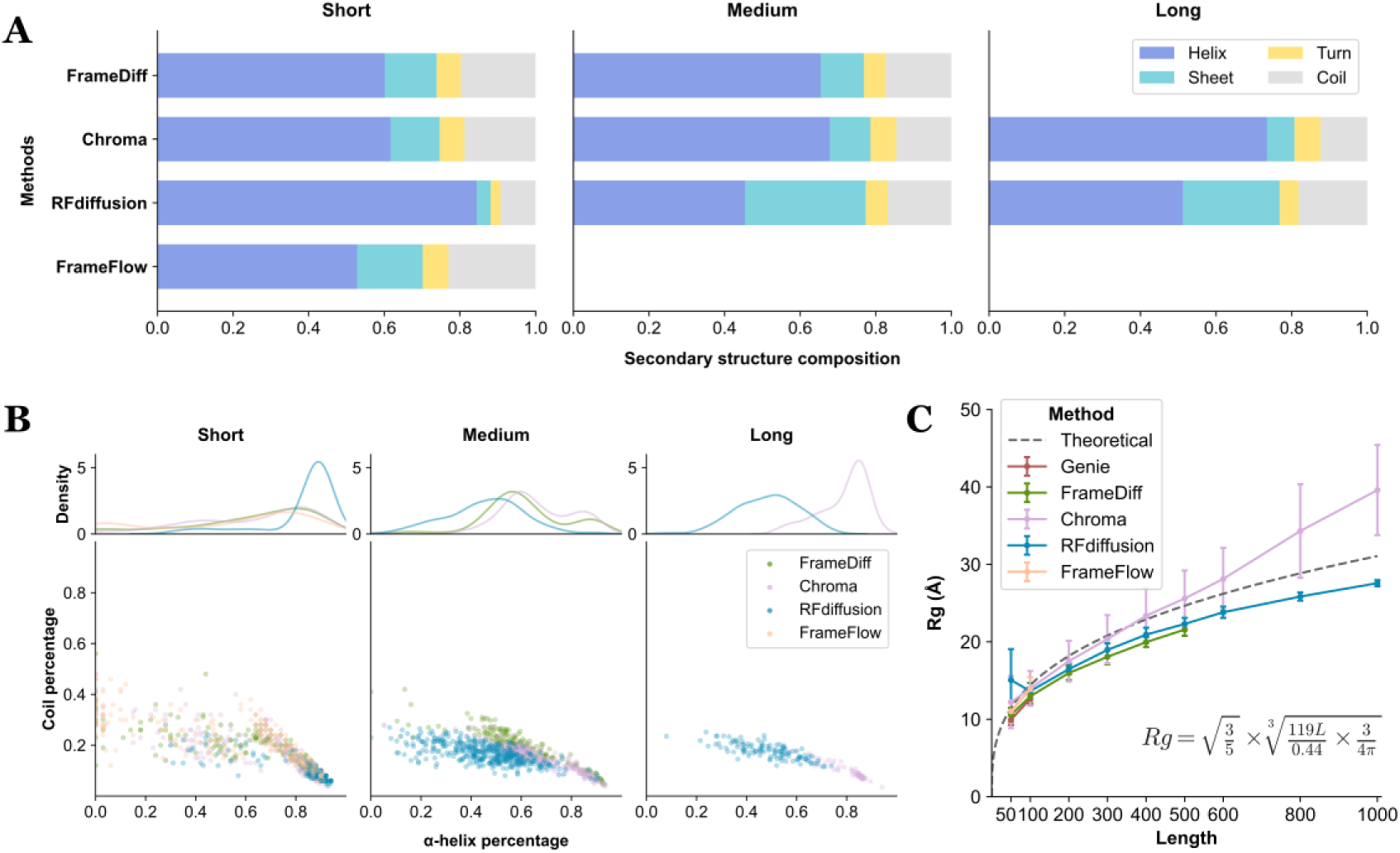
Structural analysis on generated proteins. **(A)** Average secondary structure distribution within each method. **(B)** Distribution of secondary structure composition of each protein. All data was curated by directly calculating onto protein backbones without refolding. *Genie* was excluded from this test as it solely generates *C⍺*-only backbones. **(C)** Radius of gyration (Rg) values of different methods. The points are the mean value of Rg and the error bars denote the standard deviation interval within each group. The empirical regression curve of Rg for structured proteins is shown in a dashed style with the text denoted the formula based on previous studies^73,74^.

As the overall average value of SSEs might not reflect the secondary structure of proteins faithfully at an individual level, we also analyzed the density distribution of SSEs of individual proteins, whose results are shown in **Figure 3B**. The distribution of short protein group is relatively broader than the others. *RFdiffusion* exhibits a strong tendency to generate proteins consisting mostly of *⍺*-helices, while it tends to produce proteins with more diverse SSEs than *Chroma* as length increases. This distribution combined with the overall suggest that the methods tend to generate helical proteins, which is reasonable not only for the abundant presence of *⍺*-helices but also potentially because the short-range interactions are easier for models to learn compared with the long-range interactions in *β*-sheets and flexible interactions in coils.

#### Radius of gyration

Apart from secondary structures, Rg was also calculated due to its importance to reveal the compactness of protein, whose results are shown in **Figure 3C**. With the empirical regression curve of Rg for structured proteins as a reference, it is shown that most of the methods tends to generate proteins with Rg below the value of corresponding lengths, exhibiting the compactness of generated proteins. *Chroma* is the only one exception with more stretched backbones with more lengths, indicating a potential to design more flexible protein structures.

### Ranking of Different Methods

After the analysis of generated protein backbones, we use a ranking procedure based on a procedure combined with TOPSIS decision-making and CRITIC weighting-selection technique, whose result is gathered in **Table 3**. We ranked the methods based on the length grouping of short, medium and long, respectively. Based on the ranking outcome, *FrameFlow* and *Genie* performs the best on short proteins, while *RFdiffusion* shows an outstanding performance upon generating long proteins. We also noticed that no single method consistently performs best or worst across all groups, reflecting significant different in the scenarios each method excels in. Consequently, we recommend a flexible and personalized selection of methods in practical applications, tailored to achieve the most optimal performance under specific circumstances.

**Table 3.**
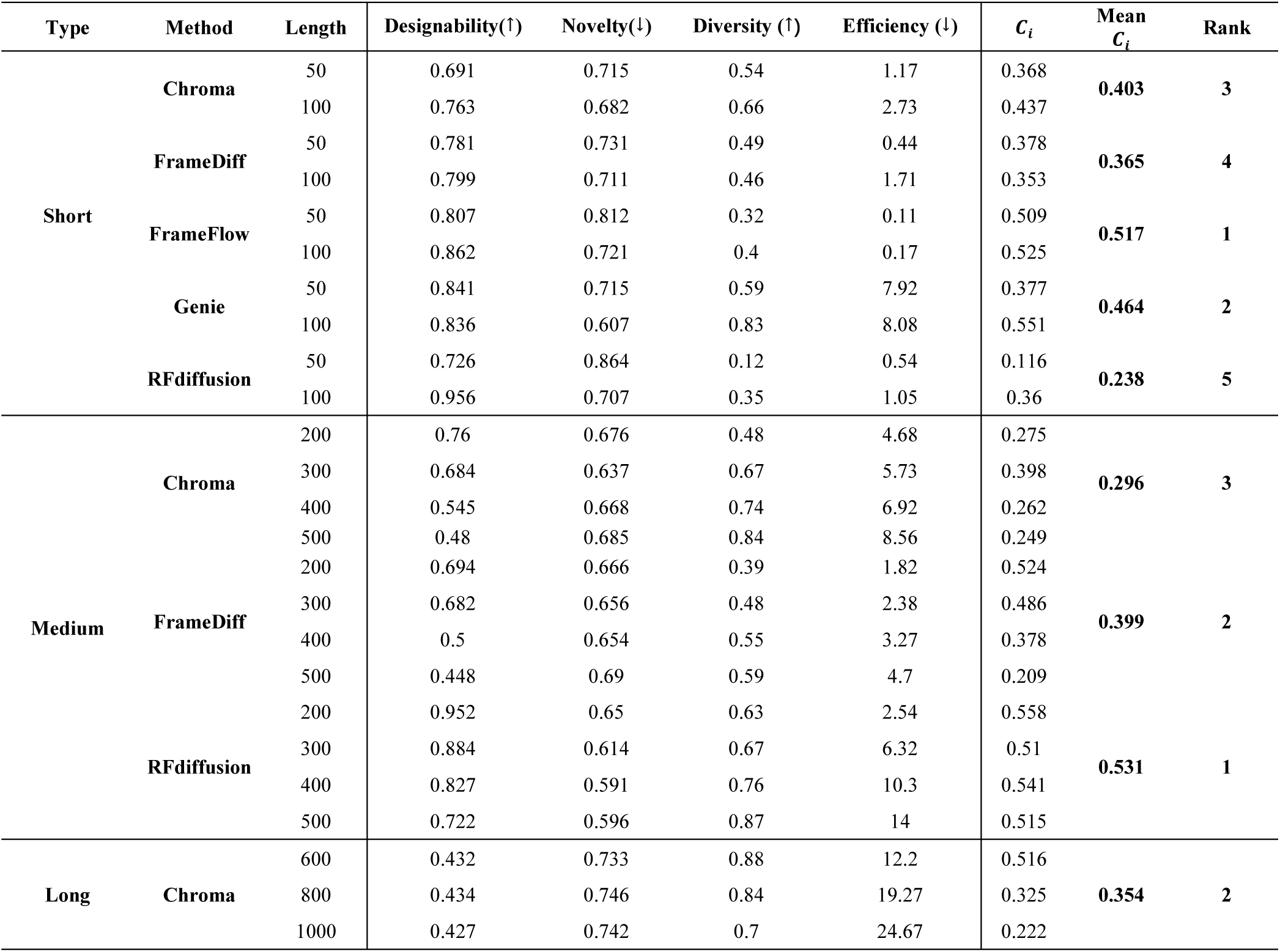
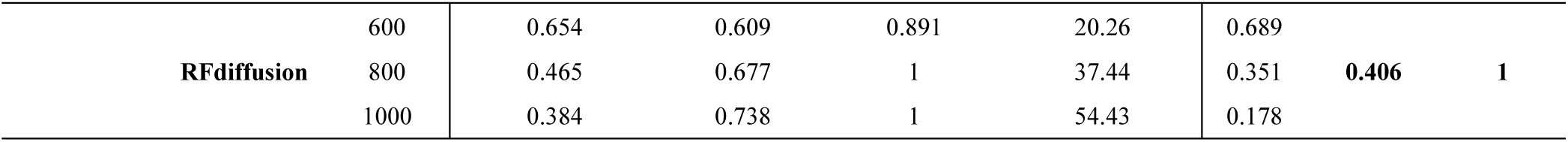
Ranking of Different Methods. Four metrics with arrows are shown as mean values within each group without data preprocessing. *C*_*i*_ denotes the calculated relative nearness value of TOPSIS results, where higher indicates better performances.

### Conditional Generation

Apart from the more conceptual unconditional generation, we extended our analysis to motif-scaffolding as a representative problem of conditional generation. We conducted a comprehensive experiment across the 24 cases described above. Similar to the unconditional task, we defined a success backbone with refolding structures a sc-TM > 0.5 and motif-RMSD < 1.0 Å, with the number of sequences selected by “top *N*” strategy, where we set *N* = 3 in this case.

**Figure 4A** displays the average performance on designability of four tested methods along 24 cases. We observed that *RFdiffusion* and *GPDL* exhibited strong efficacies with an average success rate nearby 40%, during which *GPDL* and *TDS* generated considerably diverse backbones. All methods showcased high variances of distribution, implicating that their performances varied among different tasks. To explore the differences of success rate and diversity between distinct cases, we computed these metrics for each individual case (shown in **Supplementary Figure 6**), where results show that different methods have their particular strengths in designing certain cases. We inspected that the cases can be loosely divided to three categories: *Simple*, *Moderate* and *Difficult*. *Simple* cases can be designed quite well along all tested method, such as *2KL8* (**Figure 4B**), while *Moderate* cases show diverging performances by different methods such as *7MRX_60* (**Figure 4C)**. *Difficult* cases, like *4JHW* **(Figure 4D)**, prove challenging to all methods. We argue that this diverging performance potentially result from different secondary structures and topologies of certain motifs as well as the length scopes of motifs and scaffolds for different cases. For instance, tackling *2KL8* needs designing a 20-residue scaffold around a 59-residue fixed motif, where the predefined well-structuralized motif allows models to sculpt a small scaffold more easily. To make a comparison between different success definition, we also calculated the individual success rate under the definition using sc-RMSD < 2.0 Å instead of sc-TM > 0.5 (shown in **Supplementary Figure 7**), which showcases a similar trend.

**Figure 4.**
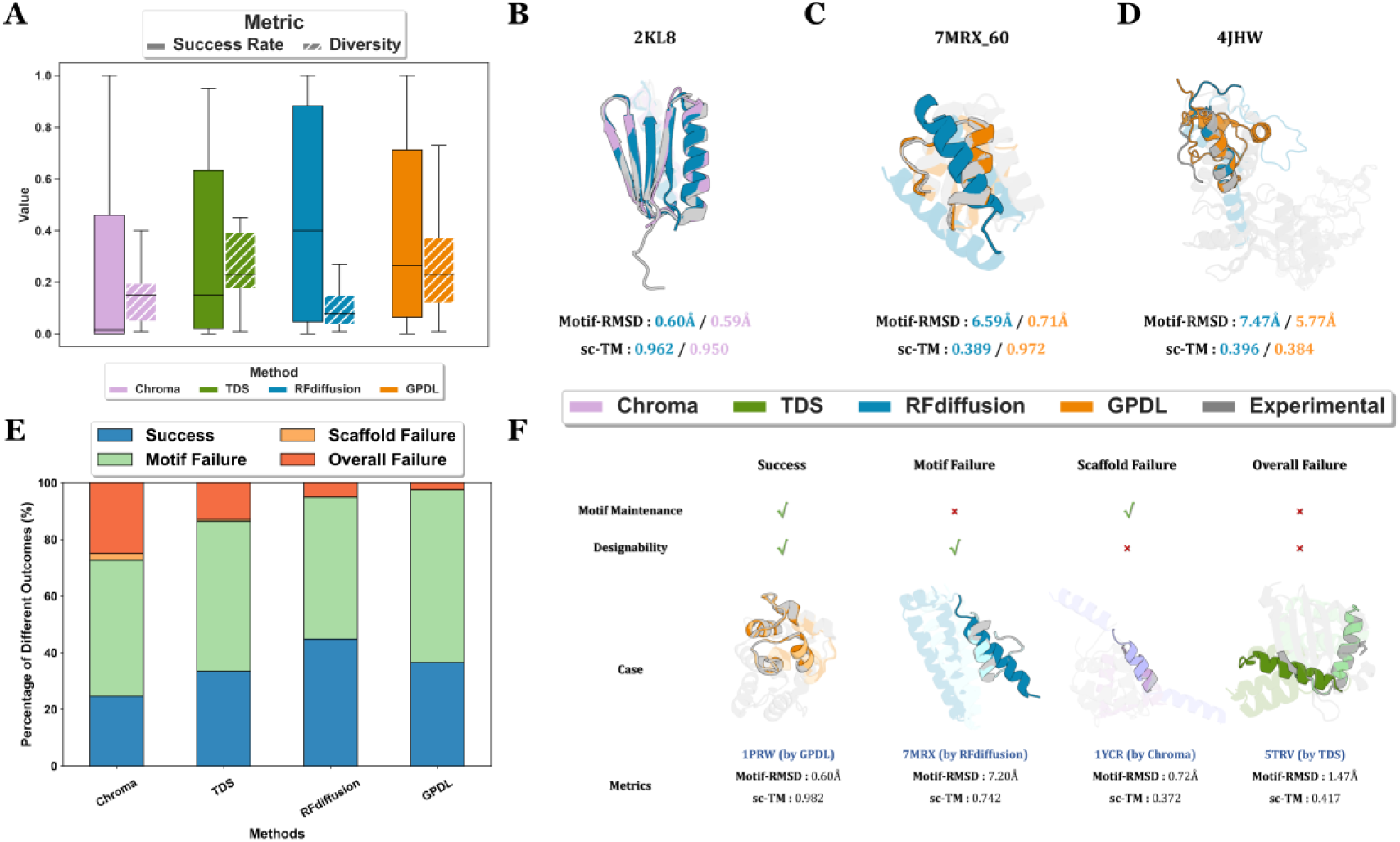
Motif-scaffolding results and failure mode analysis for tested methods. **(A)** Overall distribution of success rate and diversity across the 24-cases benchmark. The success definition follows the “Top N” strategy in unconditional generation. **(B)**, **(C)**, **(D)** The difficulty of solving motif-scaffolding problems varies across different cases. **(B)** An illustration of *Simple* cases represented by *2KL8*. **(C)** An illustration of *Moderate* cases represented by *7MRX_60*. **(D)** An illustration of *Difficult* cases represented by *4JHW*. In **(B)**, **(C)** and **(D)**, refolded structures of designs by different methods (shown in the legend on top of Figure 4F) are overlayed with crystal structures from PDB (shown in gray). Motifs are displayed as *opaque* and scaffolds are displayed as *transparent*. **(E)** Percentage of different outcomes of motif-scaffolding task among the 24-cases dataset. Each color denotes a unique type of outcome. **(F)** Schematic diagram of different outcomes. Each case in the “case” panel denotes a unique type of outcome and different colors denote different methods (shown in the legend on top of Figure 4F). Inside each scheme, we use *dark* colors to represent *refolded structures* after folding and *light* colors to denote *original backbones* designed by different methods. Experimental structures are shown in gray, motifs are displayed as *opaque* and scaffolds are displayed as *transparent*. The motif-RMSD is calculated between refolded structures and experimental structures from PDB.

We intended to gain a deeper understanding on the potential causes of unsuccessful designs beyond the explicit performances of different methods. Thus, we performed an analysis on the different outcomes of each motif-scaffolding task by grouping them into four categories. Since the success criterion by satisfying both motif-RMSD < 1.0 Å and sc-TM > 0.5 corresponds separately to motif maintenance and overall designability, we further characterized the failure cases into three types: We defined the designs that fulfill motif maintenance but not overall designability as *motif failure* and the opposite ones as *scaffold failure*, plus the ones that failed to satisfy either criterion as *overall failure*. A failure mode analysis was conducted based on this classification which is shown in **Figure 4E**. It could be easily observed that *motif failure* dominated the failed cases, followed by *overall failure*. It is noteworthy that *scaffold failure* accounts for a minimal fraction of failed cases, indicating that meeting the overall designability is a necessary but not sufficient condition for a successful design.

The analysis on failure modes offers us a chance to dive deeper into the nature of conditional generation. The fact that overall designability is a prerequisite for solving the motif-scaffolding problems emphasizes the necessity of learning a reasonable protein structure compared to meeting a certain substructure. Since the motif-scaffolding problem is a representative task of conditional generation, we propose that this mode can be generalized to the broader domain of conditional generation. In other words, designing a physically-plausible protein structure is more essential than fulfilling certain conditions. Based on this concept, we supposed that extending a well-established unconditional generation model to conditional generation by certain modifications or fine-tuning is more efficient than developing a conditional generation model from scratch, as the former strategy stems from a model that has already captured the underlying distributions of protein structures to a certain extent.

## Discussion

In this work, we conducted a thorough study of different protein backbone generation methods across various types of tasks, providing a precise and comprehensive evaluation. Beyond the qualitative rankings and comparative analyses, it is worthwhile to acknowledge that diversity in short proteins and designability in long proteins represent two primary challenges across different methods. Conversely, given that *novelty* is beside the point to practical biological applications and the lack of substantial disparities it exhibited among the methods, we suggest that its significance may have been overstated in prior studies.

Designing extremely short and large proteins holds particularly utility in specific contexts, such as the former conductive for some peptide drugs^75^ or mini-binders^76,77^ and the latter for large multi-domain proteins^78^ or symmetrical protein assemblies^79,80^. While a number of peptide^81^ and binder design^82^ methods have been proposed to address these needs, the design of large proteins remains an enduring challenge. Moreover, methods focused on generating protein complexes or dynamic backbones, which are key to the design of enzymes or allosteric proteins, are still in a nascent stage.

As for the case of conditional generation, we suggest from the results that this field is still in an early era where there remains a gap to bridge before reaching practical downstream applications. The fact that *motif failure* accounts the most of failures indicates that meeting certain conditions are still challenging for these models. While several sequence design methods have been developed to tackle the task of conditional generation^83,84^, protein backbone generation methods still hold considerable potential for enhancement. The direct modeling of protein structures stands a powerful approach to manipulate the functions and desired properties of designed proteins. We propose that both clearer definitions of numerous conditional generation problems and customized training strategies like amortization would be instrumental in improving the capacities of these methods. Despite the exceptional capability of powerful generated models, the considerable performance of *GPDL* demonstrates that model-based methods like^85–87^ still hold relevance in this field. Considering the high-throughput virtual backbone generation demand for downstream tasks, the improvement of efficiency is another promising avenue for exploration.

Given that our evaluation showcases the performance of the current outstanding methods, it also paves the way for future directions in this field. The design targeting long proteins is still in a nascent stage which urgently requires enhancement. The analysis on structural properties reveals a strong preference in structured and helices-abundant proteins, preventing them to generate much coil regions and intrinsically disordered proteins. These proteins are currently studied through molecular dynamics (MD) simulation^88–90^ or conformational sampling^91^, which are discarded from most hand-picked training sets of protein backbone generation methods aiming to improve the fitness of models. An example of these tasks is the design of non-idealized proteins attempting to deal with the aforementioned preferences of current methods, which has seen progress in ^92^. We suggest that a higher-level representation for protein structures as well as augmentation in training data would be highly useful for the generalizability of different methods. With the rapid advancement in recent years, we are witnessing an avalanche of sophisticated tasks in protein backbone generation coming into view, offering greater challenges and prospect in protein discoveries.

## Acknowledgements

We would like to thank Xiaochen Cui, Xiaoyue Ji and Taeyoung Choi for insightful comments. We thank Tao Guo for helpful discussions on practical usage related to Foldseek-Cluster. This work was supported by the Center for HPC at Shanghai Jiao Tong University, and the National Key Research and Development Program of China (2023YFF1205102 and 2020YFA0907700), the Fundamental Research Funds for the Central Universities (YG2023LC03), and the National Natural Science Foundation of China (21977068 and 32171242).

