## Supplementary Material for "Scaffold-Lab: Critical Evaluation and Ranking of Protein Backbone Generation Methods in A Unified Framework"

**Hai-Feng Chen (Full Professor)**

State Key Laboratory of Microbial metabolism, Joint International Research

Laboratory of Metabolic & Developmental Sciences, Department of Bioinformatics

and Biostatistics, National Experimental Teaching Center for Life Sciences and

Biotechnology, School of Life Sciences and Biotechnology, Shanghai Jiao Tong

University, Shanghai, 200240, China

**Notes**

The authors declare that there is no conflict of interest.

**Author Contributions**

† These authors contributed equally to this work.

**Evaluation Metrics**

**Designability**

The designability of protein backbones is the most vital metric for protein backbone generation tasks, as it directly determines the utility and reliability of generated protein backbones. The designability of a protein was originally defined as whether there exists a protein sequence that could be accommodated by a given protein backbone structure, represents the sequence-structure mapping relationship[11, 12]. It was initially studied on physical-based lattice models that utilized a reduced-alphabet consisting solely of hydrophobic and non-hydrophobic amino acids, namely *HP models* [13-15]. This approach was then employed to estimate how many feasible number of amino acid sequences the given structure could accommodate. These early studies already indicated two important aspects of the assessment of protein designability: The probability of finding amino-acid sequences that can fold into the protein’s three-dimensional structure, and the number of these optimal sequences. Despite of the relative simplicity of these models, they have indicated the variations in the capacity to accommodate sequences and the designability among different proteins, laying a crucial conceptual foundation for protein design and protein structure prediction.

As a huge number of sequences can be found given a protein backbone, it is rather unrealistic to evaluate the designability of the protein backbones through wet-lab validation. Fortunately, with the emergence of highly accurate protein structure prediction methods in recent years[3, 5, 10, 16-18], the improvement on both the precision and computational efficiency allows researchers to build an evaluation pipeline for protein backbone generation methods at a good pace. The first holistic evaluation process in this field, to our best knowledge, was raised by *ProtDiff* [19], which conceptualize the self-consistency TM-score[20] (sc-TM) by calculating the TM-score between the original generated backbone and the predicted structure of sequence designed from the backbone. Since proteins with a mutual TM-score greater than 0.5 are widely considered to belong to the same fold pattern[21], the sc-TM can, to a certain extent, reflect the mapping relationship between sequences identified by the backbone and the backbone itself. This evaluation process comprises three critical components: the *backbone generator* that generates protein backbones, and the *sequence designer* that generates sequences for protein backbones, and the *structure predictor* that predict structures for generated sequences, is often referred as the “refolding” pipeline, where the three components organize three in a row and has already become a paradigmatic way for the evaluation protein backbones.

Based on the current progress, we aim to establish a comprehensive experiment to conduct a systematic and thorough performance evaluation of the selected methods. We used ProteinMPNN[22] for its robust performance shown in wet-lab experiments[8, 23] and ESMFold[10] for the sake of computational cost and its better performance on *de novo* proteins compared with AlphaFold2[24]. Building upon the existed refolding pipeline, we further expanded the evaluation process along the original definition of designability by incorporating the robustness of protein backbones. We additionally adapted two modifications:

1. We used sc-TM instead of sc-RMSD throughout our evaluation pipeline. Although many works used a threshold of sc-RMSD < 2.0 Å to define the success of backbones, we found this metric showcased commonly high values and hard for scaling during preliminary experiments, while sc-TM showed a greater robustness. Nevertheless, we calculated the designability and success rate using sc-RMSD for a comparison and the integrity of our evaluation, which can be found in **Supplementary Figure 3-4**.
2. “Top $N$ strategy”. While current works considered a backbone as success when there is solely one sequence satisfy the criteria, we argue that the number of sequences that can fold into the generated structures reveal the robustness and fitness of backbones to some extent. So during our experiment, we reported the sc-TM as the mean value of $N$ sequences with highest sc-TM among generated sequences, where the formula is defined as follows:

$$Success_{backbone}=\left\{ \begin{aligned} 1,\frac{1}{N}\sum_{i=1}^{N} scTM>0.5 | (scTM_{1}>\ldots>scTM_{N}>scTM_{i}) \\ 0, others \end{aligned} \right.$$

During our evaluation, we chose $N=3$, which means we calculated the designability use the mean value of top 3 sequences with highest sc-TM values out of 10 selected sequences. We also reported the distribution of sc-TM and success rate by picking the best sequence as a fair comparison, which are displayed in **Figure 2B** and **Supplementary Figure 5**.

Furthermore, as it was found that more sequences generated by ProteinMPNN might lead to a higher success rate, which is reasonable for the extension of sequence searching space. In this evaluation pipeline, we aim to develop a way not only take a step to the vast sequence searching space but also maintain the computational cost, so we design 100 sequences using ProteinMPNN for each generated protein backbone and select 10 sequences with the lowest negative log likelihood, which is a parameter already been used to select sequences generated by ProteinMPNN and proved to have certain positive relationship with the stability of sequence itself.

For the *success rate* of motif-scaffolding problem, we followed the one used in *RFdiffusion*, except replacing the AlphaFold2 single-chain version with ESMFold for the sake of computational efficiency and comparable prediction accuracy between them. For the backbone part we take ESMFold sc-TM < 0.5 and ESMFold pAE < 5 Å, while for the motif part we take ESM motif-RMSD < 1.0 Å, which we referred as “backbone designability” and “motif designability”, respectively. The overall success rate are defined by finding sequences that both satisfying the two aspects of designability, while the number of sequences is carried on as the “Top $N$” strategy we used in the task of unconditional generation, while in this section we also choose$N = 3.$ We also noticed that a new success definition on motif-scaffolding task by treating motifs and scaffolds separately shas been brought out recently in [25], whereas we didn’t conduct this due to the scope of our work, but as a constructive metric to be studied in the future.

The predicted local-distance difference test (pLDDT) and predicted aligned error (pAE) raised by AlphaFold2 are two candidates for the backbone success definition in our evaluation pipeline. While they have potential in measuring designability and the latter one was once used for success definition in *RFdiffusion*, we argue they are not the best choice in our evaluation for two particular factors: First, they are highly relative to particular protein structure prediction methods, which might brought external bias to the pipeline; Second, we found a strong relationship between sc-TM and these two metrics respectively during our preliminary experiments, whose results we display in **Supplementary Figure 8**. Nevertheless, they are still potential useful under some specific circumstances, such as pLDDT for special regions in proteins and pAE for multi-domain proteins. We suggest using them independently for special cases.

**Novelty**

When considering the task of protein backbone generation, especially in the case of *de novo* protein design, we noticed that the ultimate objective of protein design is to design non-existed proteins, in other words, proteins with novel properties compared with those existed in nature. Since the three-dimensional structure of protein has a strong correlation with its functions, it is necessary to assess how similar these newly generated backbones to existed proteins in nature. Besides its downstream applications, the novelty of generated protein backbones is also an important metric for models themselves because it implies the out-of-distribution generalizability of model, since most of the methods use native proteins datasets like CATH[26, 27] and Protein Data Bank (PDB)[28, 29] as training set.

Some of the previously methods have already conducted experiments on the novelty analysis of generated protein backbones, with most of them focusing on finding the most similar protein structure in large protein databases like PDB, to the generated protein backbones. Next a similarity metric is carried out by calculating the TM-Score between these two structures, which is often referred as “pdbTM”. If this value is low, it means that the generated backbones are novel compared to most of the existing experimentally solved native proteins, and vice versa.

We consider pdbTM a valuable metric for calculating novelty of generated protein backbones. Nevertheless, different methods use different searching tools and searching parameters, resulting in a gap to compare the novelty of different methods fairly. In this work, we use Foldseek[30] to search generated backbones against PDB database to find their nearest protein structure. In order to make a fair comparison while maintaining computational efficiency, we apply the same parameter suite to all tested methods.

Beyond the original definition of pdbTM, we found a phenomenon that though some of the protein backbones has low pdbTM values, they also behave bad in the aspect of designability, which is reasonable since some of the unrealistic protein backbones can hardly find similar protein partners in PDB naturally. Thus, we argue that some modifications should be taken to maintain the balance between novelty and designability. The way we take is to add a penalty to those with low designability, where the definition is formulated as follows:

$$S_{novelty}=\left\{ \begin{aligned} pdbTM+\left( 0.5-w\frac{1}{n}\sum_{i=1}^{n} scTM \right),\frac{1}{n}\sum_{i=1}^{n} scTM<0.5 \\ pdbTM,\frac{1}{n}\sum_{i=1}^{n} scTM>0.5 \end{aligned} \right.$$

Here we define a novelty score by putting a penalty on less-designable backbones.

**Diversity**

We utilized Foldseek-cluster to cluster different groups and calculated the diversity value as then. Though different combination set of parameters had been tried, we found Foldseek-cluster failed to cluster certain groups where error occurred during the prefilter step. We hypothesis this was because some of the groups has a relatively low diversity that Foldseek-cluster could not assign enough non-redundant secondary structures. We used TM-score threshold = 0.5 during clustering throughout the evaluations process, and additional results by using TM-score threshold = 0.4 and TM-score threshold = 0.6 are displayed in **Supplementary Figure 9**. Generally, two proteins with a TM-score above the TM-score threshold are tend to be assigned to the same cluster, so the diversity might get higher as the value of threshold increases.

**Ranking Methods**

Throughout our ranking procedure, we used a combination of CRITIC technique for weight determination of different indicators and TOPSIS technique for the normalization and scoring of different methods.

**CRITIC**

The CRiteria Importance Through Intercriteria Correlation (CRITIC) is a technique that obtains the objective weights of relative importance which is widely utilized by decision-makers. It uses the standard deviation $S_{j}$ to indicate the variation of values within each indicator. A larger standard deviation indicates a greater variation in protein design methods, which reflects more information and stronger evaluation intensity for that indicator, for which it is assigned a higher weight relative to others.

Consider we have $i$ different methods evaluated by $j$ different indicators, the mean value  $\bar{x}_{j}$ and standard deviation $S_{j}$ of each indicator is formulated as below:

$\left\{ \begin{aligned} \bar{x}_{j}=\frac{1}{n}\sum_{i=1}^{n} x_{ij} \\ S_{j}=\sqrt{\frac{{\sum_{i=1}^{n} (x_{ij}-\bar{x}_{j})}^{2}}{n-1}} \end{aligned} \right.$

Where $n$ is the total number of evaluated methods.

After that, we calculate the correlation between different indicators, $R_{j}$, to assign the weights of them accordingly. For each indicator, the higher it correlates to other indicators, the lower the conflicts between it and other indicators:

$$R_{j}=\sum_{i=1}^{P} (1-r_{ij})$$

Where $r_{ij}$ is the correlation parameter between the evaluation indicators $i$ and $j$ and $P$ is the number of indicators. Generally, a higher value of $R_{j}$ indicates a greater similarity in the information reflected and a higher degree of redundancy in the evaluation context. To some extent, this weakened the evaluation strength of the indicator and the weight given to it should be reduced.

Next, the amount of information, $I_{j}$, is calculated accordingly:

$I_{j}=S_{j}\sum_{i=1}^{P} (1-r_{ij})=S_{j}\times R_{j}$

For each indicator $j$, a large value of $I_{j}$ implies its crucial role within the evaluation, which is assigned a high weight relative to those with a small $I_{j}$.

Based on the above, the objective weight $w_{j}$ of the indicator $j$ is determined as follows:

$w_{j}=\frac{I_{j}}{\sum_{j-1}^{P} I_{j}}$

Then we can obtain the objective weight of each indicator $W^{1}$.

**TOPSIS**

TOPSIS (Technique for Order Preference by Similarity to Ideal Solution) technique is a multi-criteria decision analysis method which based on the concept that the chosen alternative should have the shortest geometric distance from the positive ideal solution (PIS, $S^{+}$) and the longest geometric distance from the negative ideal solution (NIS, $S^{-})$. Compared to the non-compensatory methods that eliminate alternatives that do not meet a particular criterion, TOPSIS as a representative of compensatory methods allows trade-offs between criteria, where a poor result in one criterion can be negated by a good result in another criterion, which provides a more realistic form of modeling of data.

To calculate $S^{+}$ and $S^{-}$ for TOPSIS, we use the weight matrix $W$ obtained by a weight assignment method, which is CRITIC in our study. We multiply the matrix$X$ by $W$ to obtain the matrix $V$, as shown below:

$$V=\left[ \begin{matrix} V_{11} & V_{12} & \ldots& V_{1n} \\ V_{21} & V_{22} & \ldots& V_{2n} \\ \ldots& \ldots& \ldots& \ldots\\ V_{m1} & V_{m2} & \ldots& V_{mn} \end{matrix} \right]=\left[ \begin{matrix} X_{11} & X_{12} & \ldots& X_{1n} \\ X_{21} & X_{22} & \ldots& X_{2n} \\ \ldots& \ldots& \ldots& \ldots\\ X_{m1} & X_{m2} & \ldots& X_{mn} \end{matrix} \right]\times\left[ \begin{matrix} W_{1} & 0 & 0 & 0 \\ 0 & W_{2} & 0 & 0 \\ 0 & 0 & \ldots& 0 \\ 0 & 0 & 0 & W_{n} \end{matrix} \right]$$

At this point

$\left\{ \begin{aligned} S^{+}=max\left\{ V_{ij} | 1\leq i\leq m \right\} \\ S^{-}=min\left\{ V_{ij} | 1\leq i\leq m \right\} \end{aligned} \right.$

The geometric distance to PIS and NIS is calculated respectively as below:

$\left\{ \begin{aligned} D_{i}^{+}=\sqrt{\sum_{j=1}^{n} {(V_{ij}-S^{+})}^{2}} \\ D_{i}^{-}=\sqrt{\sum_{j=1}^{n} {(V_{ij}-S^{-})}^{2}} \end{aligned} \right.$

The relative nearness $C_{i}$ of each solution is then formulated:

$C_{i}=\frac{D_{i}^{-}}{D_{i}^{-}+D_{i}^{+}}$

Where $C_{i}$ is the final parameter we used for ranking in **Table 3** in main text. Based on $C_{i}$, we rank the solutions where larger values indicate better design methods.

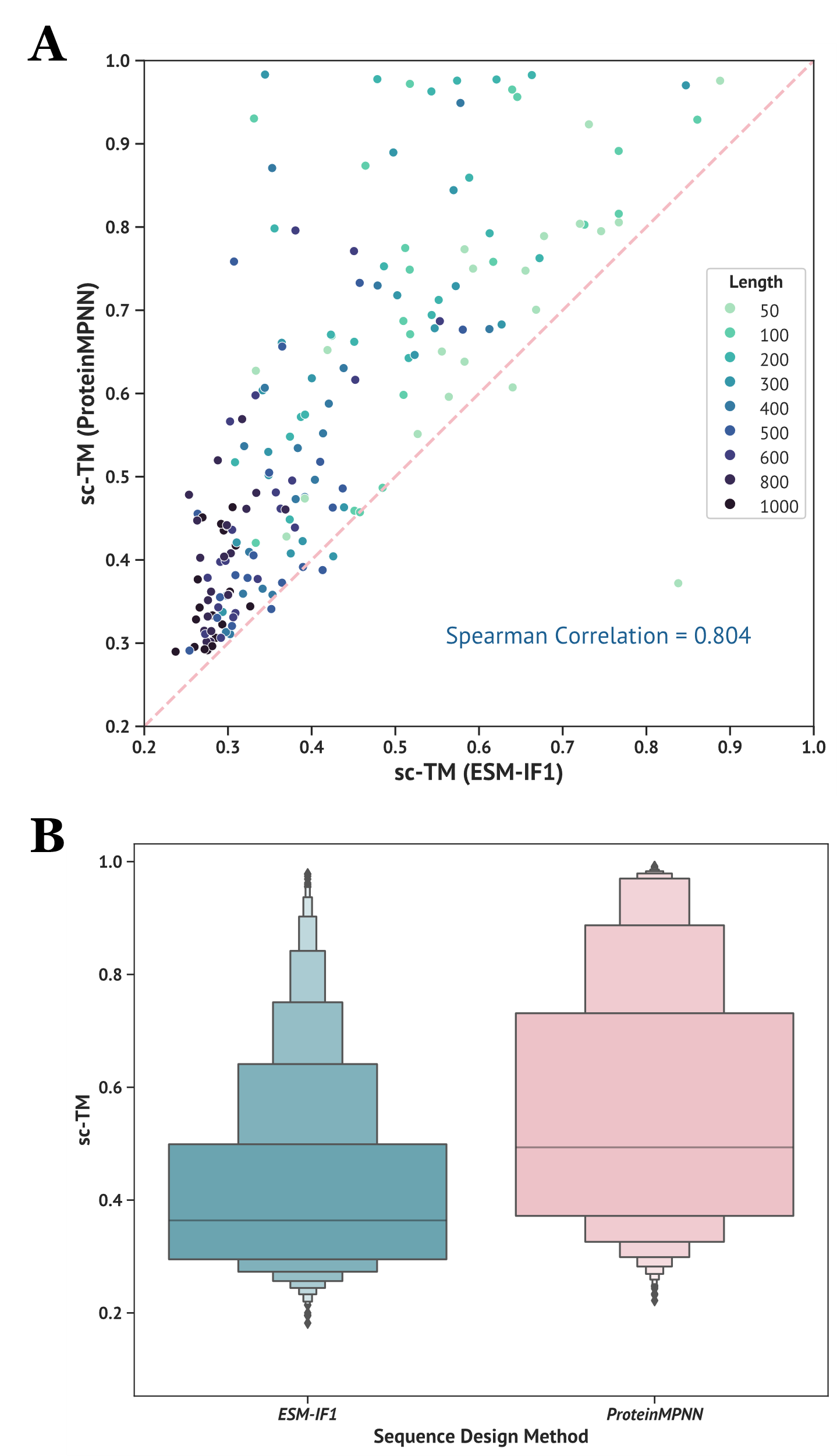

**Supplementary Figure 1. Compared studies on different sequence design methods.** To make a comparison between ProteinMPNN and ESM-IF1, we randomly selected 20 backbones generated for each tested length (from 50 to 1000) and designed 10 sequences for each backbone, resulting in a subset containing 9 (number of unique lengths) × 20 = 180 unique backbones and 180 × 10 = 1800 designed sequences. We then feed these sequences into ESMFold and calculated the values of self-consistency TM-score (sc-TM). **(A)** Correlation between ProteinMPNN and ESM-IF1. Each point in the plot denotes a unique backbone, $x$ and $y$ axis denote the sc-TM values origined from ProteinMPNN and ESM-IF1, respectively. The degree of color denotes the number of residues of each sequence. The value displayed on the lower right denotes the Spearman correlation between results from different sequence design methods. **(B)** Distribution of sc-TM values from ProteinMPNN and ESM-IF1 are displayed in a boxenplot. It can be observed that sequences obtained from ProteinMPNN exist significant higher sc-TM values than ESM-IF1 no matter from the perspective of single backbones or overall distribution.

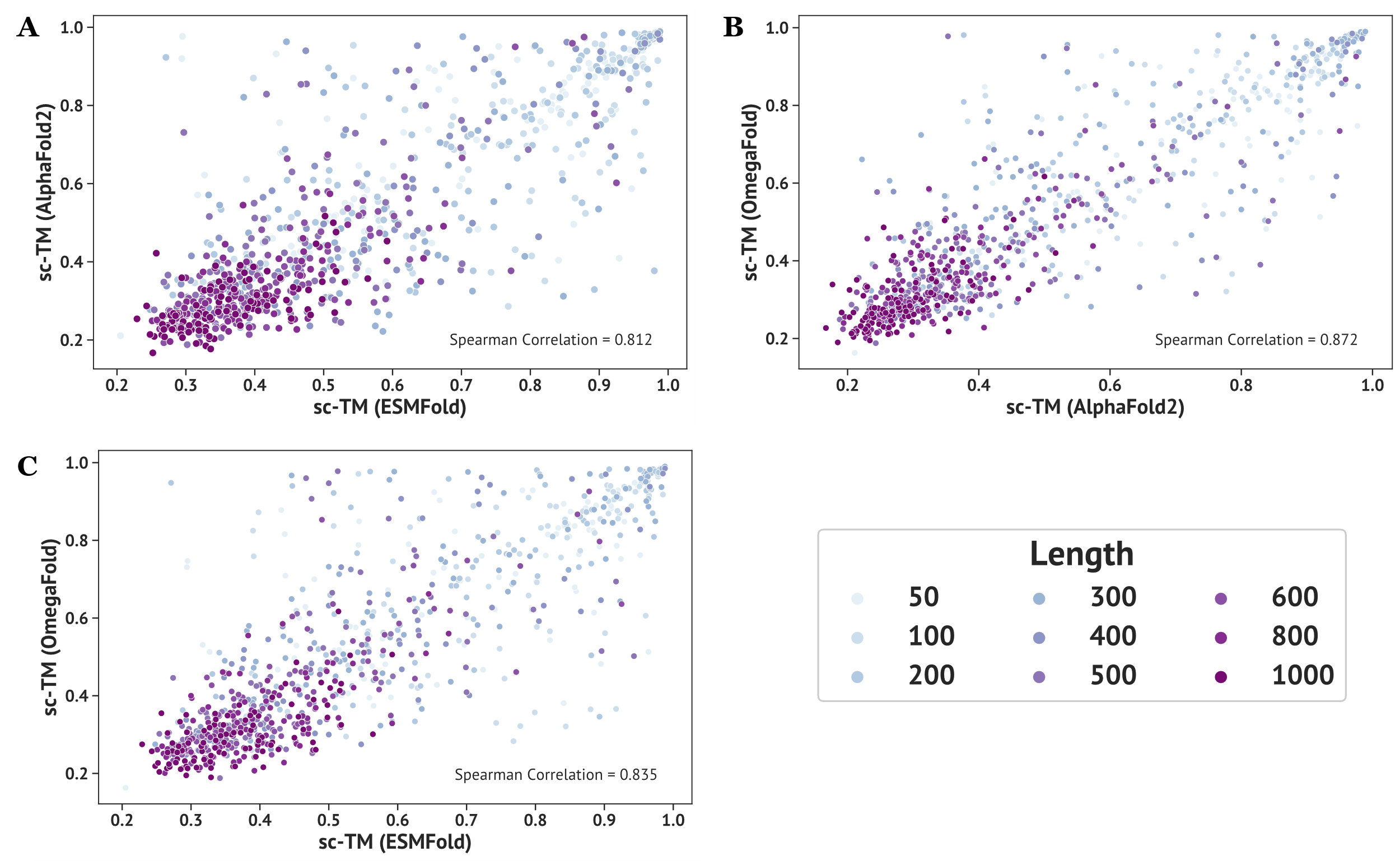

**Supplementary Figure 2. Correlation between three protein structure prediction methods.** For comparison, we randomly selected sequences designed by ProteinMPNN from generated backbones and fed them into three structure predictors. For each length of unconditional sampled backbones, we randomly selected 100 sequences, which resulted in a total number of 900 sequences predicted by ESMFold, AlphaFold2 (without MSA) and OmegaFold, respectively. Each point in the subplot denotes a unique sequence and the value displayed on the lower right denotes the Spearman correlation between results from different structure predictors. The degree of color denotes the number of residues of each sequence. **(A)** Correlation between ESMFold and AlphaFold2 (without MSA). **(B)** Correlation between AlphaFold2 (no MSA) and OmegaFold. **(C)** Correlation between ESMFold and OmegaFold.

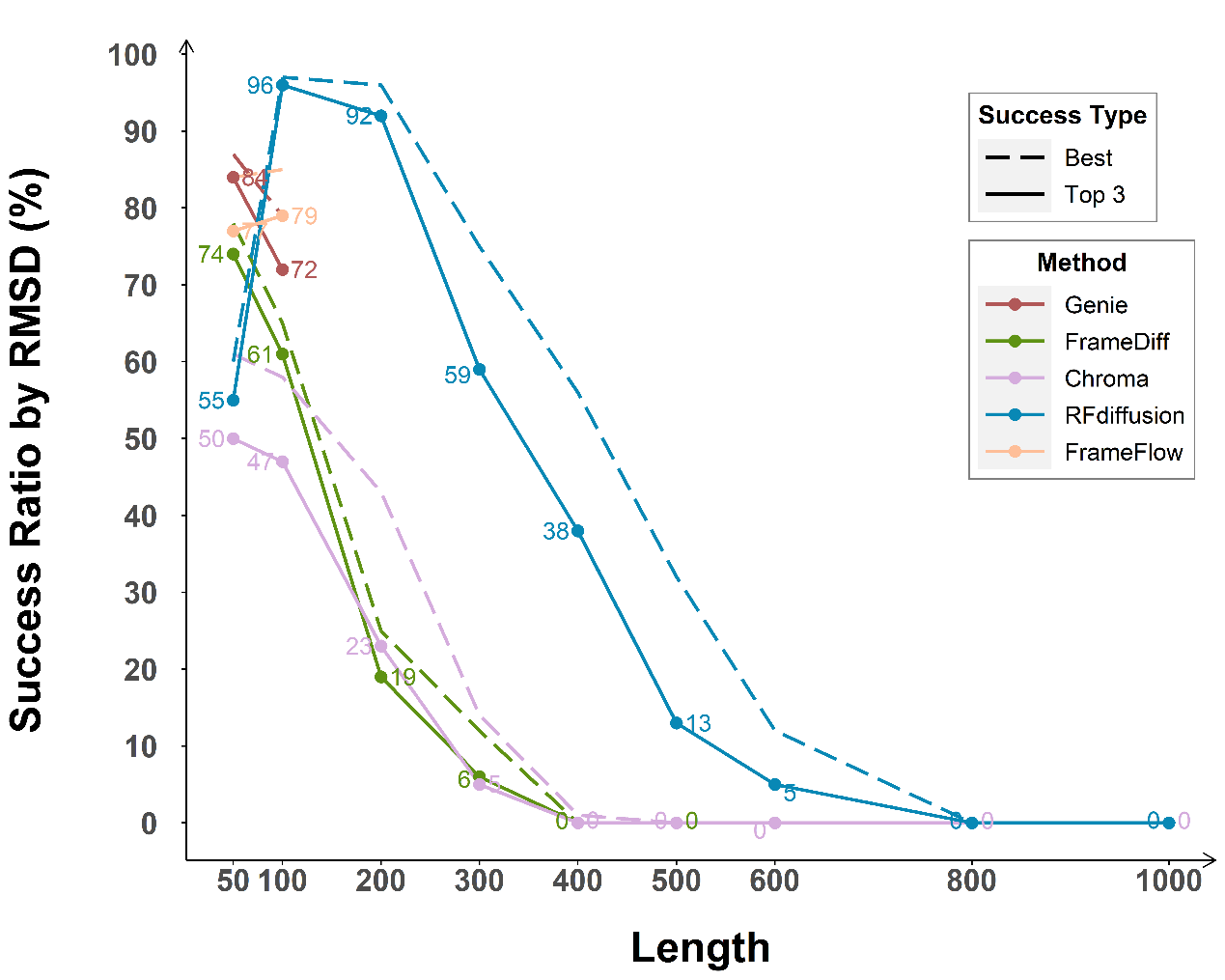

**Supplementary Figure 3. Success rate defined by sc-RMSD for different methods.** As a comparison, the success rate here is defined by the number of backbones passing the sc-RMSD threshold (top 3 /best sc-RMSD < 2.0 Å) divided by the total number of backbones within each group, with top 3 sc-RMSD by solid line and best sc-RMSD by dashed line. The text of numbers refers to the solid lines.

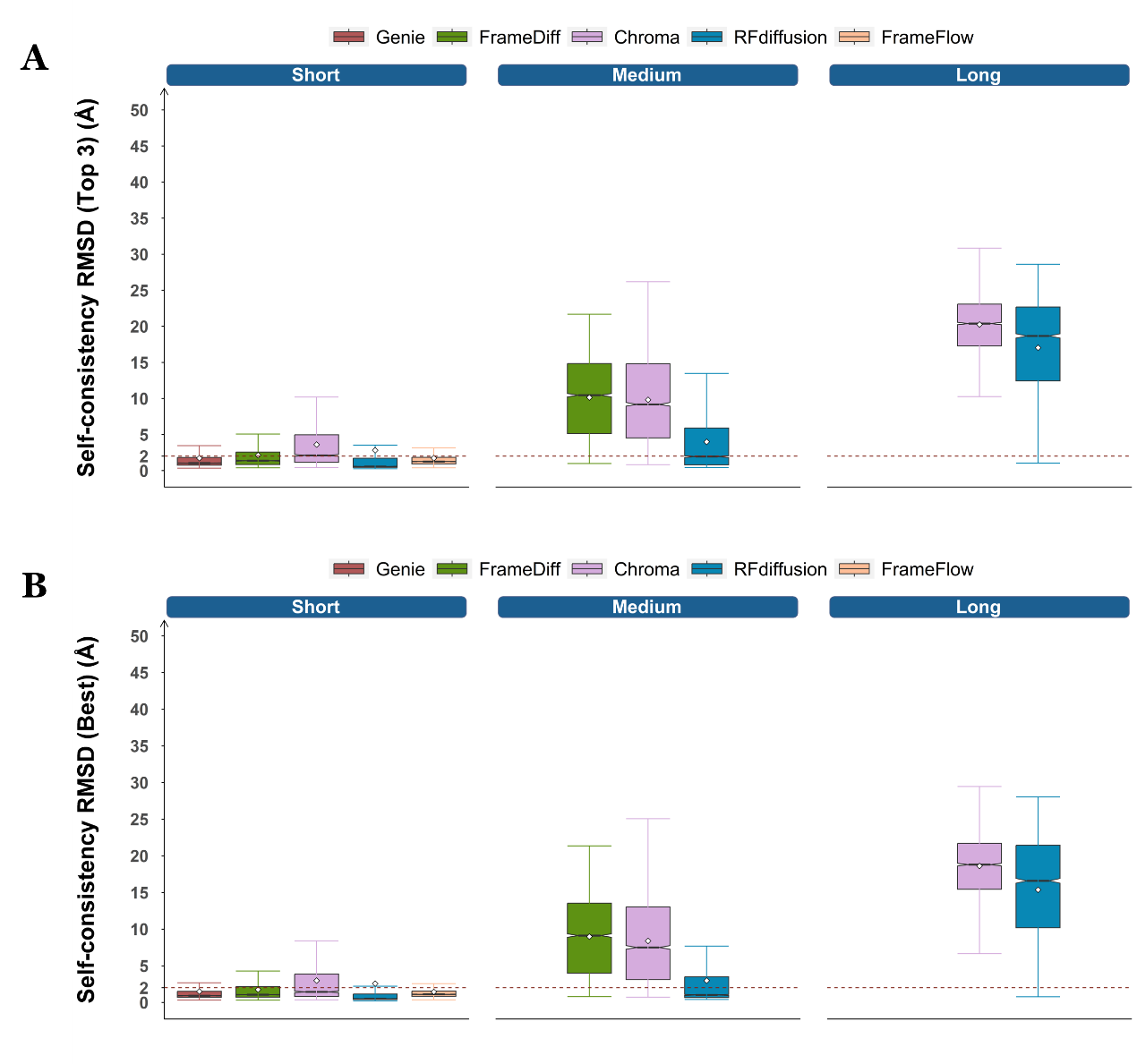

**Supplementary Figure 4. Distribution of top 3 / best sc-RMSD for different methods.** The white diamonds denote the median value of each group and the dashed line shows the designability threshold of sc-RMSD = 2.0 Å. **(A)** Distribution of top 3 sc-RMSD. **(B)** Distribution of best sc-RMSD.

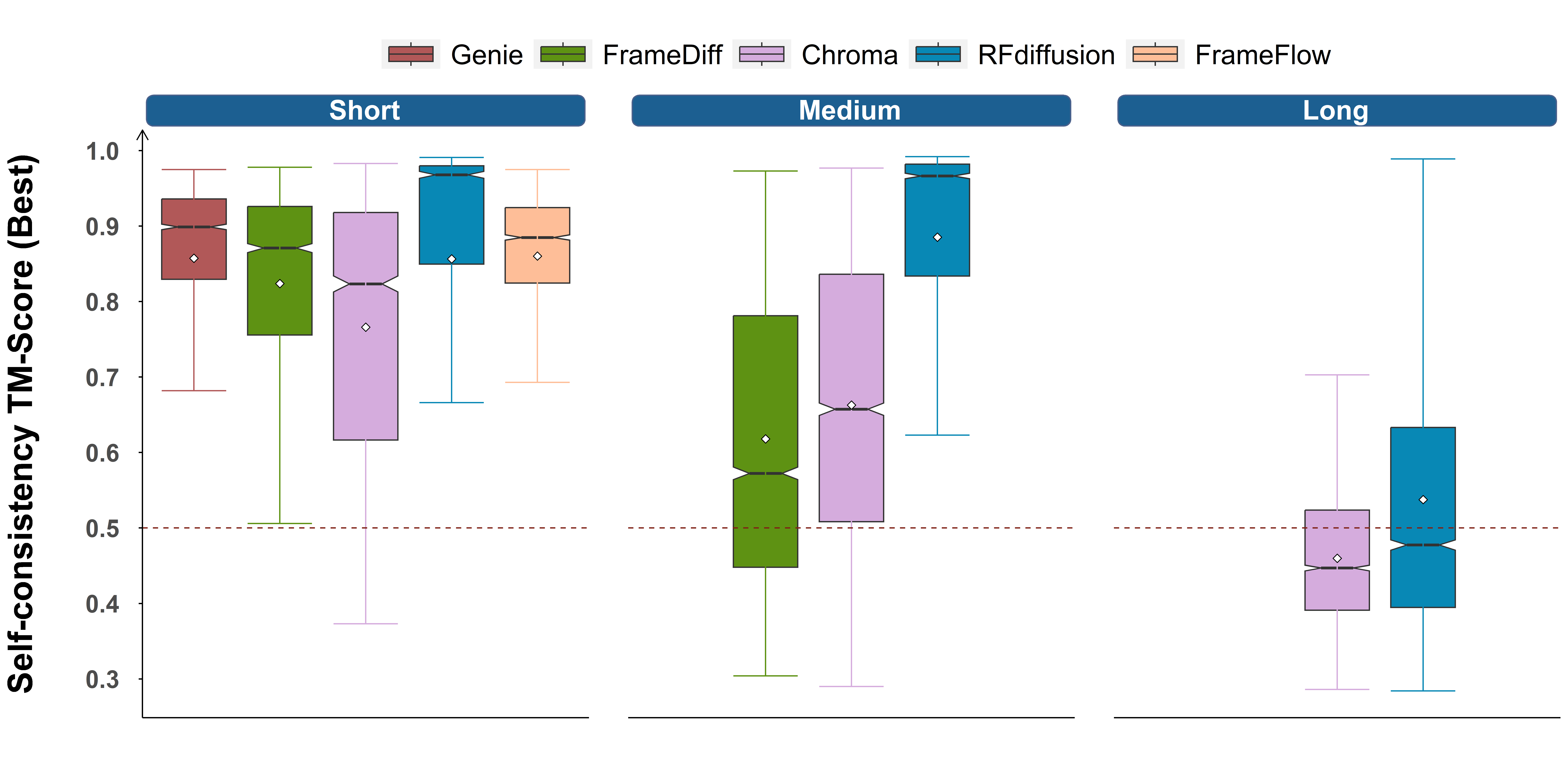

**Supplementary Figure 5. Distribution of best sc-TM for different methods.** The white diamonds denote the median value of each group and the dashed line shows the designability threshold of sc-TM = 0.5.

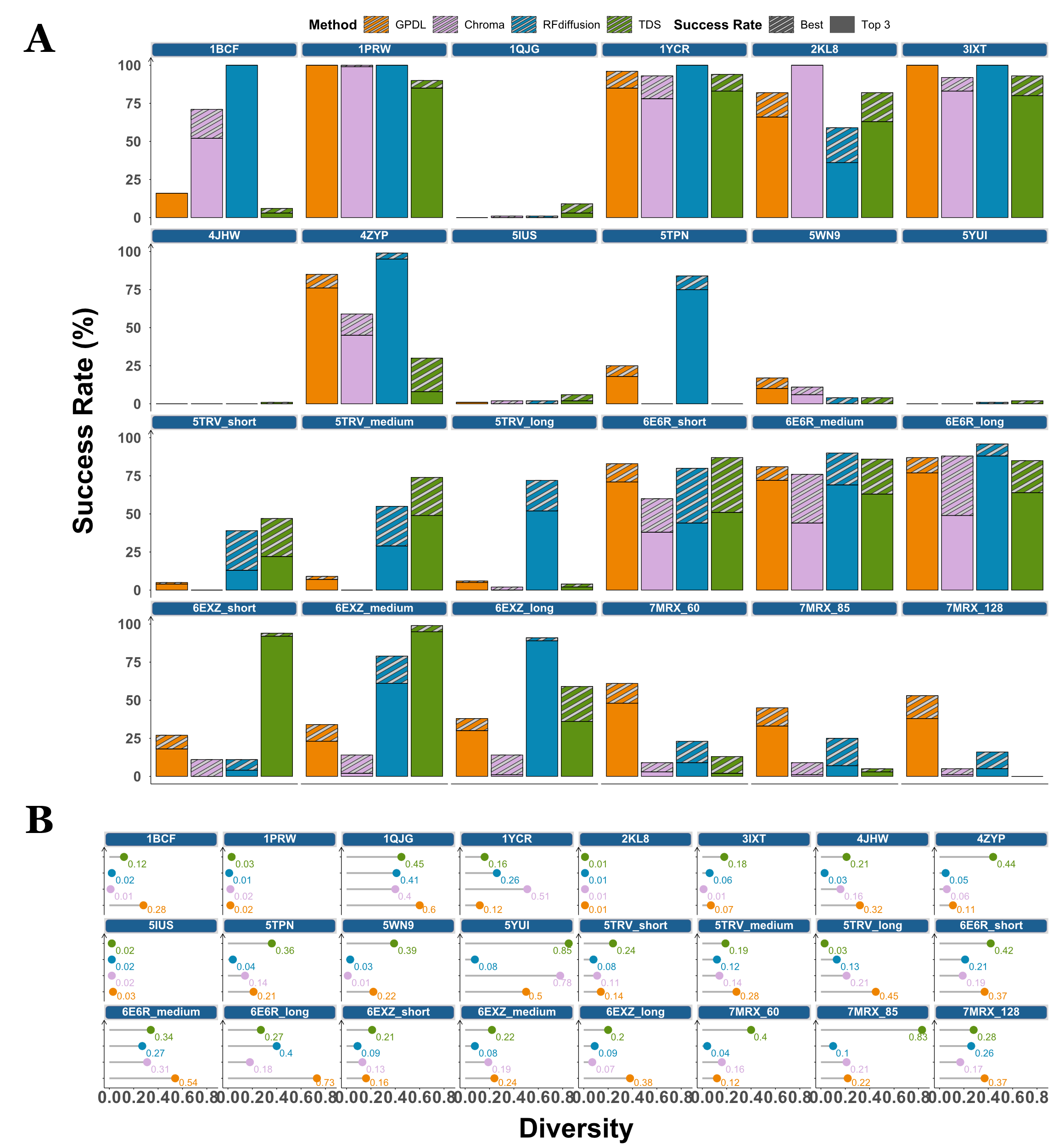

**Supplementary Figure 6. Detailed Motif-scaffolding performances for tested methods.** Each facet places the result of one specific cases of four methods. **(A)** Success Rates of different methods on designability. The success threshold is defined as refolded sc-TM of the whole structure > 0.5 and motif-RMSD < 1.0Å. The definition of success rate follows the ones with unconditional generation, with the areas with pure color denote the top 3 success rates and the shadow areas as the value surpassed by best success rates. **(B)** Diversity of different methods. The lollipops display the diversity values of sdifferent methods.

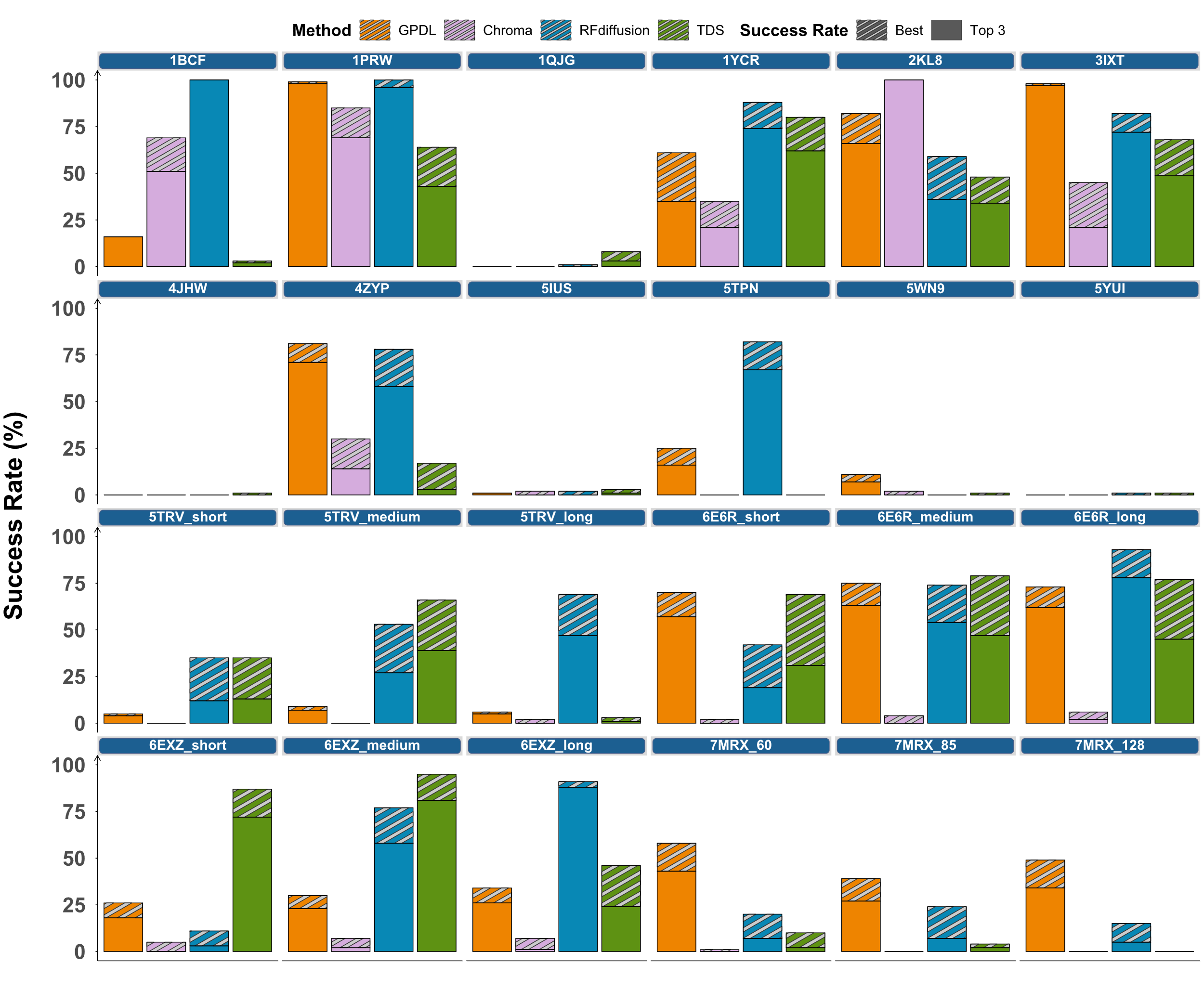

**Supplementary Figure 7. Motif-scaffolding performances using backbone success as sc-RMSD < 2.0** **Å.** Each facet places the result of one specific cases of four methods. The success definition follows the ones in **Figure 4** except changing the backbone success by sc-RMSD. The overall success threshold is defined as refolded sc-RMSD of the whole structure < 2.0 Å and motif-RMSD < 1.0 Å. The areas with pure color denote the top 3 success rates and the shadow areas as the value surpassed by best success rates.

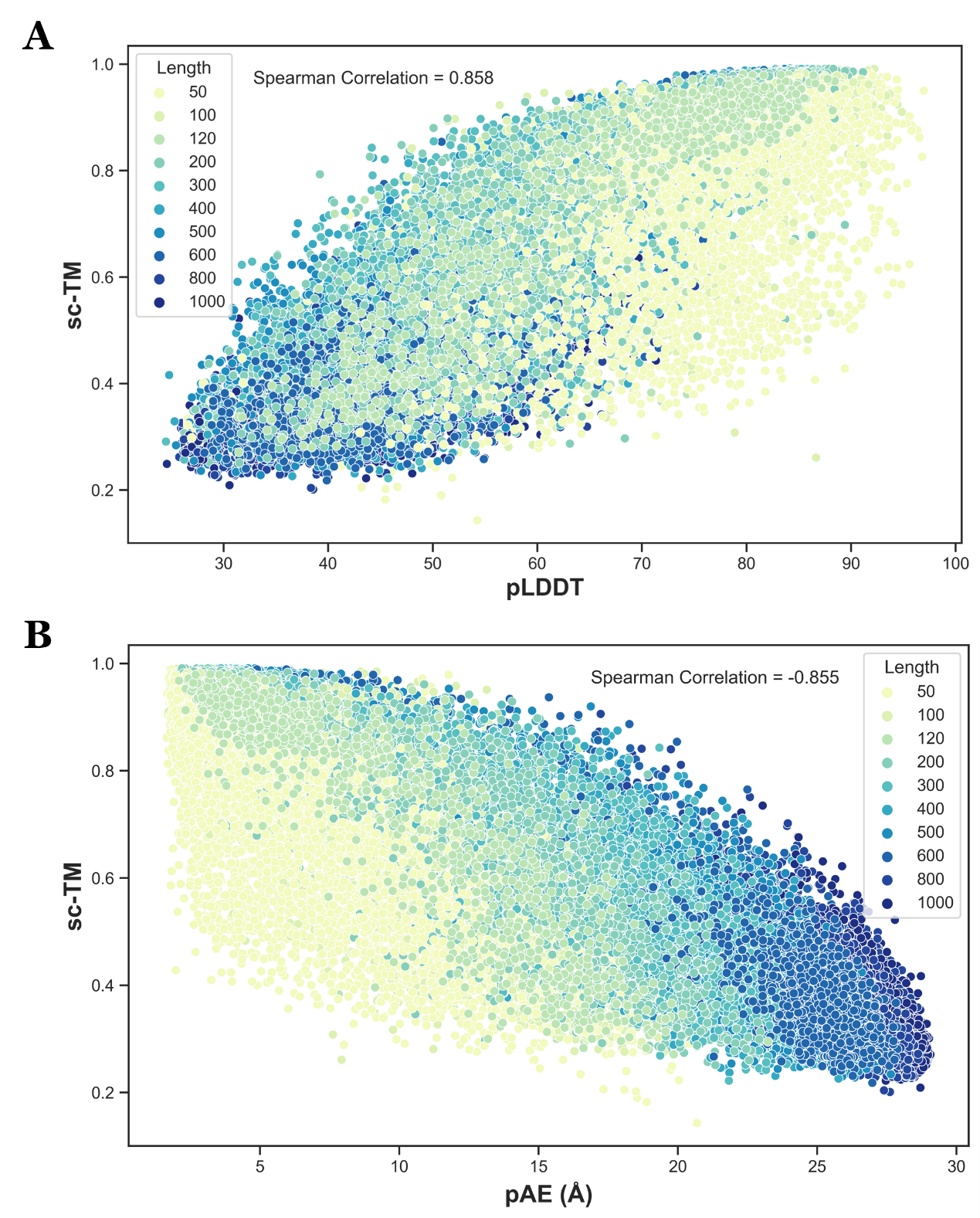

**Supplementary Figure 8. Correlation between different metrics.** The points displayed here are all protein backbones generated in the task of unconditional generation, where different colors denote the length of protein. The Spearman correlation value between two metrics is displayed on the plot. **(A)** Correlation between ESMFold pLDDT and self-consistency TM-score. **(B)** Correlation between ESMFold pAE and self-consistency TM-score.

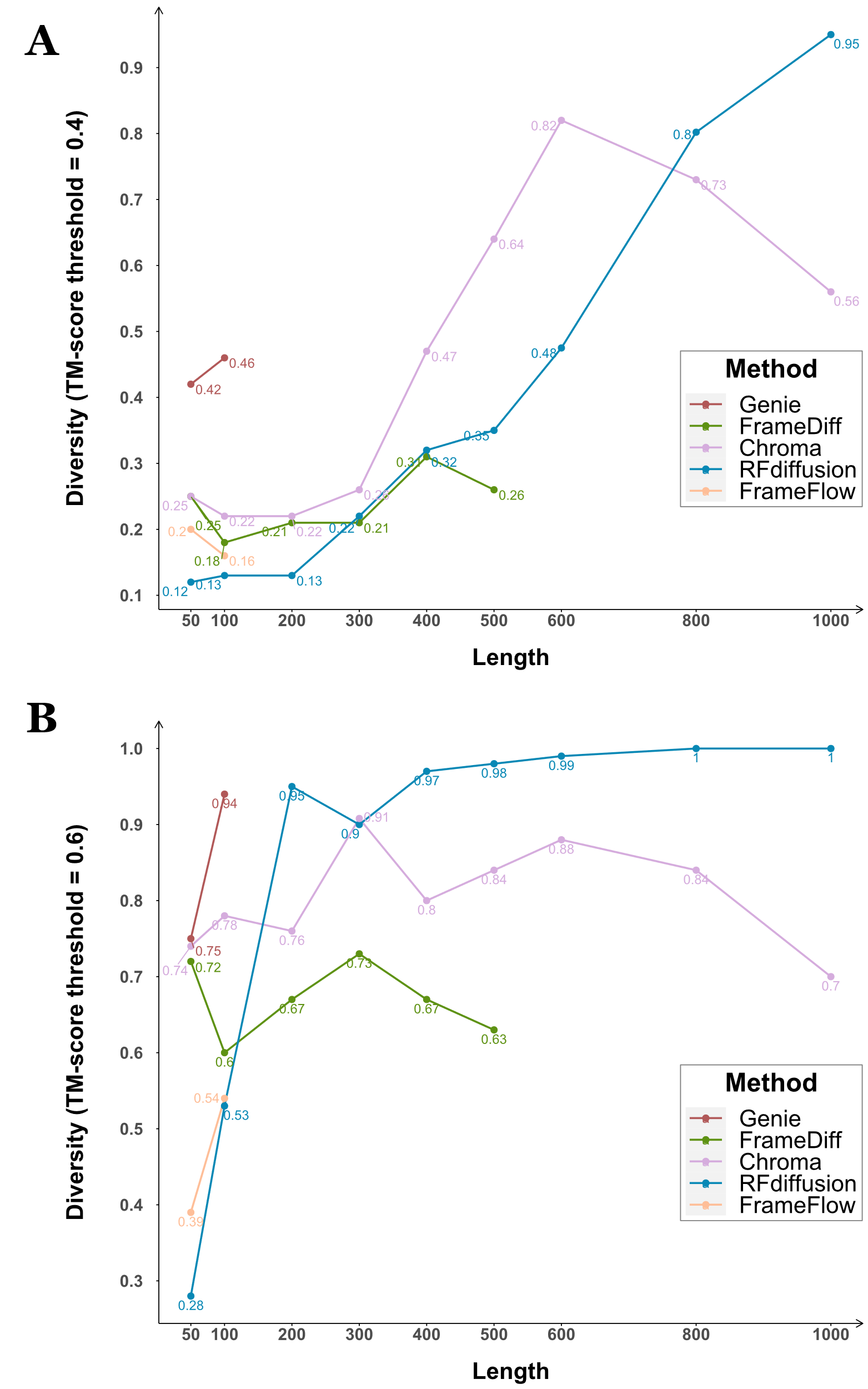

**Supplementary Figure 9. Diversity results for unconditional generation using different TM-score threshold by Foldseek-Cluster.** **(A)** Diversity values of TM-score threshold = 0.4. **(B)** Diversity values of TM-score threshold = 0.6.

**Supplementary Table 1. 24 cases selected for evaluation on motif-scaffolding.**

| **Case (PDB ID)** | **Input** | **Total Length (*RFdiffusion & Chroma*)** | **Total Length (*TDS*)** | **Total Length (*GPDL*)** |
| --- | --- | --- | --- | --- |
| ***1PRW*** | 5-20/**A16-35**/10-25/**A52-71**/5-20 | 60-105 | 100 | 80 |
| ***1BCF*** | 8-15/**A92-99**/16-30/**A123-130**/16-30/**A47-54**/16-30/**A18-25**/8-15 | 96-152 | 125 | 112/115 |
| ***5TPN*** | 10-40/**A163-181**/10-40 | 50-75 | 75 | 69 |
| ***5IUS*** | 0-30/**A119-140**/15-40/**A63-82**/ 0-30 | 57-142 | 100 | 107 |
| ***3IXT*** | 10-40/**P254-277**/10-40 | 50-75 | 75 | 73/74 |
| ***5YUI*** | 5-30/**A93-97**/5-20/**A118-120**/10-35/**A198-200**/10-30 | 50-100 | 75 | 91 |
| ***1QJG*** | 10-20/**A38**/15-30/**A14**/15-30/**A99**/10-20 | 53-103 | 100 | 83 |
| ***1YCR*** | 10-40/**B19-27**/10-40 | 40-100 | 75 | 59 |
| ***2KL8*** | **A1-7**/20/**A28-79** | 79 | 100 | 79 |
| ***7MRX_60*** | 0-38/**B25-46**/0-38 | 60 | 75 | 60 |
| ***7MRX_85*** | 0-63/**B25-46**/0-63 | 85 | 100 | 85 |
| ***7MRX_128*** | 0-122/**B25-46**/0-122 | 128 | 125 | 128 |
| ***4JHW*** | 10-25/**F196-212**/15-30/**F63-69**/10-25 | 60-90 | 75 | 73 |
| ***4ZYP*** | 10-40/**A422-436**/10-40 | 30-50 | 50 | 45 |
| ***5WN9*** | 10-40/**A170-189**/10-40 | 35-50 | 50 | 45 |
| ***5TRV_short*** | 0-35/**A45-65**/0-35 | 56 | 50 | 56 |
| ***5TRV_medium*** | 0-65/**A45-65**/0-65 | 86 | 75 | 86 |
| ***5TRV_long*** | 0-95/**A45-65**/0-95 | 116 | 125 | 116 |
| ***6E6R_short*** | 0-35/**A23-35**/0-35 | 48 | 50 | 48 |
| ***6E6R_medium*** | 0-65/**A23-35**/0-65 | 78 | 75 | 78 |
| ***6E6R_long*** | 0-95/**A23-35**/0-95 | 108 | 100 | 108 |
| ***6EXZ_short*** | 0-35/**A28-42**/0-35 | 50 | 50 | 50 |
| ***6EXZ_medium*** | 0-65/**A28-42**/0-65 | 80 | 75 | 80 |
| ***6EXZ_long*** | 0-95/**A28-42**/0-95 | 110 | 125 | 110 |

**Case:** Different cases are denoted by their PDB IDs. We basically followed the benchmark set collected by *RFdiffusion* except ***6VW1*** since it is a complex generation task which *TDS* was not capable to perform.

**Input:** The part prefixed by a letter in bold are the inputs (chain, residues) from the **real PDB structure** provided to the model (the “functional-site” or namely the “motif”). The part without a letter prefix are the lengths that the different methods randomly sampled to generate good designs (the non-functional site or namely the “scaffold”).

**Total Length**: This indicates the total lengths of protein backbones output by different models. For *Chroma*, we performed the experiment with its SubstructureConditioner. For the other three method, we followed their original experimental settings except fixing the types of amino acids of all motif positions without redesign. The different experimental settings resulted in different sets of total lengths, where *RFdiffusion* and *Chroma* could perform length-variable design by randomly sampling from some specific length scopes, while *TDS* and *GPDL* generated proteins with mostly fixed lengths. Note that the differences between lengths of fixed or random lie on the scaffold parts, where all method were input with the same motifs from real protein structures.

**Supplementary Table 2. Significant test between whether adding the low-designability penalty during novelty calculation.** A one-sided Mann-Whitney U Test was performed within each category along different lengths and methods.

| **Type** | **Method** | **p-value** |
| --- | --- | --- |
| Short | Chroma | $0.2972$ |
|  | FrameDiff | $0.4193$ |
|  | FrameFlow | $0.4595$ |
|  | Genie | $0.4482$ |
|  | RFdiffusion | $0.4808$ |
| Medium | Chroma | $5.1874\times{10}^{-4}$ |
|  | FrameDiff | $6.1074\times{10}^{-8}$ |
|  | RFdiffusion | $0.3431$ |
| Long | Chroma | $3.3330\times{10}^{-16}$ |
|  | RFdiffusion | $2.4630\times{10}^{-15}$ |
